# Phlorotannin rich *Ascophyllum nodosum* seaweed extract inhibits influenza infection

**DOI:** 10.1101/2024.08.08.606782

**Authors:** Daniele F. Mega, Parul Sharma, Anja Kipar, Udo Hetzel, Chloe Bramwell, Alan Merritt, Samuel Wright, Chris Plummer, Richard A. Urbanowicz, James P. Stewart

## Abstract

Seaweed derived compounds are a renewable resource utilised in the manufacturing and food industry. This study focuses on an Enriched seaweed extract (ESE) isolated from *Ascophyllum nodosum.* ESE was screened for antiviral activity by plaque reduction assays against influenza A viruses (IAV) H1N1 and H3N2 subtypes. Time of addition assays and FACS analysis were used to help determine the modes of action. The therapeutic potential of ESE was then explored using differentiated human bronchiole epithelial cells at the air liquid interphase and a murine model challenged with IAV. The data indicates ESE primarily interacts directly with virions, preventing virus cell binding. Interestingly, ESE also inhibits early and late stage of the influenza A lifecycle when treatment occurs after cell binding. This inhibitory effect appears to prevent internalisation of virus and release of progeny virus by targeting neuraminidase activity. Intranasal administration of ESE in mice infected with IAV reduced viral load in lung tissue. ESE may be a promising broad acting antiviral agent in the treatment of influenza infections.

## Introduction

Influenza viruses are segmented single stranded negative sense RNA (-ssRNA) viruses in the *Orthomyxoviridae* family which are classified into 4 genera, Influenza A-D, and cause respiratory infections [1]. Influenza A viruses (IAV) are responsible for the majority of influenza infections in humans and five pandemics since 1889, the most recent of which was 2009 and the most devastating in 1918 with over 50 million recorded deaths worldwide [2], [3]. IAVs are further categorised into subtypes by their surface glycoproteins, hemagglutinin (HA; H1-H18) and neuraminidase (NA; N1-N11), of which A/H1N1 and A/H3N2 currently cause seasonal epidemics in humans [4]. Influenza viruses are constantly evolving due to genetic reassortment and low-fidelity RNA polymerase proofreading capabilities, which results in antigenic drift due to sequence changes encoding HA and NA, allowing escape from previously protective immune responses [5], [6].

There are currently several licensed antivirals and seasonal vaccines available. One such antiviral, Amantadine, is directed at the viral M2 protein, targeting viral uncoating and release of infectious nucleic acids [7], [8]. The most commonly prescribed antivirals for influenza infections are neuraminidase inhibitors (NAIs) including Zanamivir and Oseltamivir (Tamiflu), which bind to the viral NA protein, blocking enzymatic function and therefore inhibiting release of progeny virions [9], [10], [11], [12]. Resistance, however, has been reported for all currently licenced antivirals with Amantadine no longer recommended for treatment of IAV infections [13], [14], [15]. This highlights the need to develop novel broad acting antivirals, which could provide an alternative to current treatments or be used in combination with licenced antivirals.

Seaweed derived compounds have been shown to have anti-viral, anti-inflammatory and immune modulating activities [16], [17]. Fucoidans are sulphated carbohydrates found in brown algae that exhibit efficacy against IAV, binding surface glycoproteins inhibiting infection as well as the cellular EGFR pathway and viral neuraminidase activity *in vitro* [18]. Carrageenans are sulphated polysaccharides produced from red seaweed and have been well studied for antiviral activity [19]. Iota-carrageenan displays antiviral activity against rhinovirus and IAV, most likely through inhibition of cell binding or entry [20], [21].

This study focuses on the effect of a phlorotannin rich enriched seaweed extract (ESE) isolated from the brown seaweed *Ascophyllum nodosum* [22] against IAV infection *in vitro* and in a murine model.

## Results

### Anti-influenza activity of enriched seaweed extract (ESE)

The ability of ESE to inhibit virus infection in cells was screened by plaque reduction assays using three strains of IAV. Influenza A Virus A/Puerto Rico/8/1934 H1N1 (PR8), Influenza A Virus A/X-31 H3N2 (X31) and Influenza A Virus A/England/195/2009 H1N1 (Eng195) were used to represent different subtypes and a more recent pandemic strain. Non-cytotoxic concentrations of ESE were determined by measuring cell viability by MTS assay following 72 h of incubation with ESE, the longest timepoint used in the following experiments (Supplementary Figure S1). The mode of action was investigated by adding ESE one hour post infection (hpi) to the plaque assay overlay or by either pre-treating virus or cells for one hour prior to infection (Figure 1A). The number of plaques were given as a proportion of the untreated control. The pre-treatment of IAV (Figure 1D) with ESE produced the greatest reduction in plaques in a dose dependent manner, resulting in up to 100% reduction in all three strains tested at the highest concentrations. Pre-treating cells with ESE (Figure 1B) did not cause a significant reduction in plaques compared to the untreated control, suggesting ESE does not interact with cellular receptors for virus binding. The addition of ESE one hpi also resulted in a significant reduction in plaques (Figure 1C) in a dose dependent manner against all three strains and was most effective against the Eng195 strain. This data suggests non-cytotoxic concentrations of ESE most potently inhibits the early stages of the virus life cycle prior to virus cell entry while also potentially inhibiting events following cell entry.

**Figure 1.**
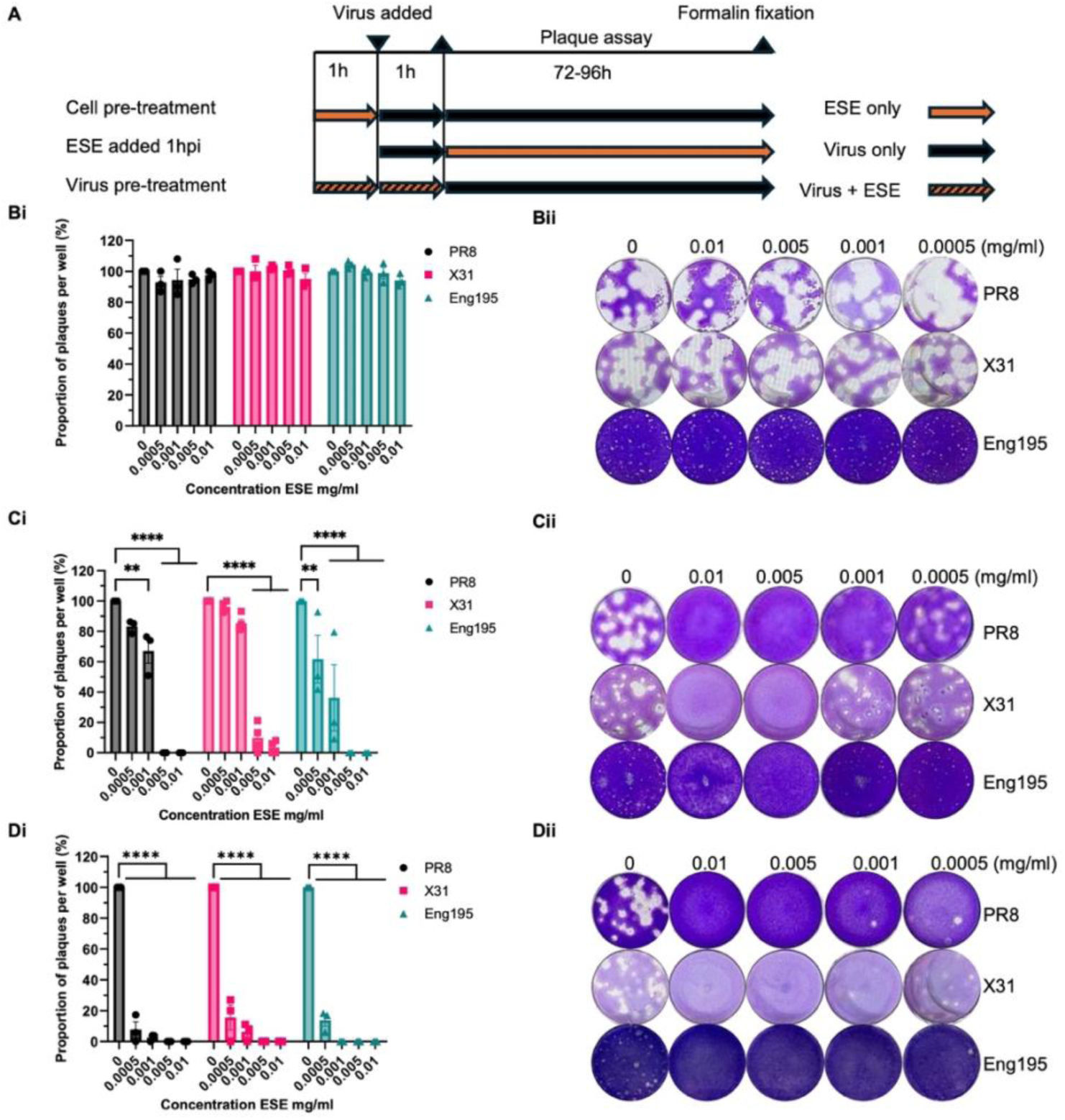
Plaque reduction assay IAV ESE. **(A)** Schematic of plaque reduction analysis. **(Bi) (Bii)** MDCK cells pre-treated with ESE or infection media. Cells were pre-treated for one hour. ESE was then removed, and cells washed with PBS prior to infection with IAV PR8, X31 or Eng195. Inoculum was removed following 1 hour of adsorption and plaque assay overlay was added until plaque formation prior to fixing and staining. **(Ci) (Cii)** ESE added following virus adsorption. MDCK monolayers were infected for one hour with IAV, inoculum was removed and plaque assay overlay containing ESE was added prior to fixing and staining upon plaque formation. **(Di) (Dii)** IAV PR8, X31 or Eng195 were pre-treated with ESE for one hour prior to infection of MDCK cell monolayers. Inoculum was removed following one hour of adsorption and plaque assay overlay was added prior to fixing and staining upon plaque formation. Data represented as mean ± SEM of three or more independent experiments. Asterisks indicate statistical difference (two-way ANOVA with Dunnett’s multiple comparisons test; **P<0.01, ****P<0.0001).

### ESE interaction with virions

The plaque reduction assays displayed greatest effect after pre-treatment of virus with ESE, suggesting a direct mode of action on the virion and inhibition of virus cell entry. This was further explored using a higher starting concentration of virus followed by serial dilutions to assess whether inhibition could be reversed or diluted out. PR8 and X31 were diluted to 10^6^ PFU/mL and mixed 1:1 with ESE for one hour prior to serial dilutions onto cell monolayers. ESE neutralised both PR8 and X31 (Figure 2A and 2B) at 0.005 and 0.01 mg/mL while 0.001 mg/mL caused approximately one log reduction in titre against both PR8 and X31. The ability of ESE to neutralise high titres of IAV despite subsequent serial dilutions suggest a strong potentially virucidal interaction between the extract and virus. The possibility of an interaction with a virus protein important for entry such as HA was investigated next. HA inhibition assay was performed for both PR8 and X31 displaying HA titres 2^5^ and 2^4^ respectively. ESE appeared to cause agglutination at high concentrations. 0.0125 mg/mL ESE inhibited agglutination at the lowest dilution of PR8 however the remaining dilutions of virus and PBS control still displayed agglutination. This may indicate ESE interacted with the high titre of virus at 2^2^ and was therefore unable to cause agglutination. 0.003125 mg/mL reduced the HA titre to 2^3^ for both PR8 and X31 while 0.0015625 mg/mL reduced HA titres to 2^4^ and 2^3^ for PR8 and X31 respectively (Figure 2C and 2D). The lowest concentration, 0.00078125 mg/mL, had no effect on HA titres for either PR8 or X31. ESE was therefore presumed to be binding viral proteins and this was analysed by separating purified IAV that had been incubated with ESE or PBS by SDS-PAGE. Coomassie staining showed viral protein bands disappeared when incubated with ESE from their expected molecular weights and appeared to aggregate at the top of the gel despite denaturing (Figure 2E), something previously seen by plant-derived tannins [23]. Treatment of BSA with ESE also caused a similar outcome, with the protein band appearing at the top of the gel following silver staining (Figure 2F).

**Figure 2.**
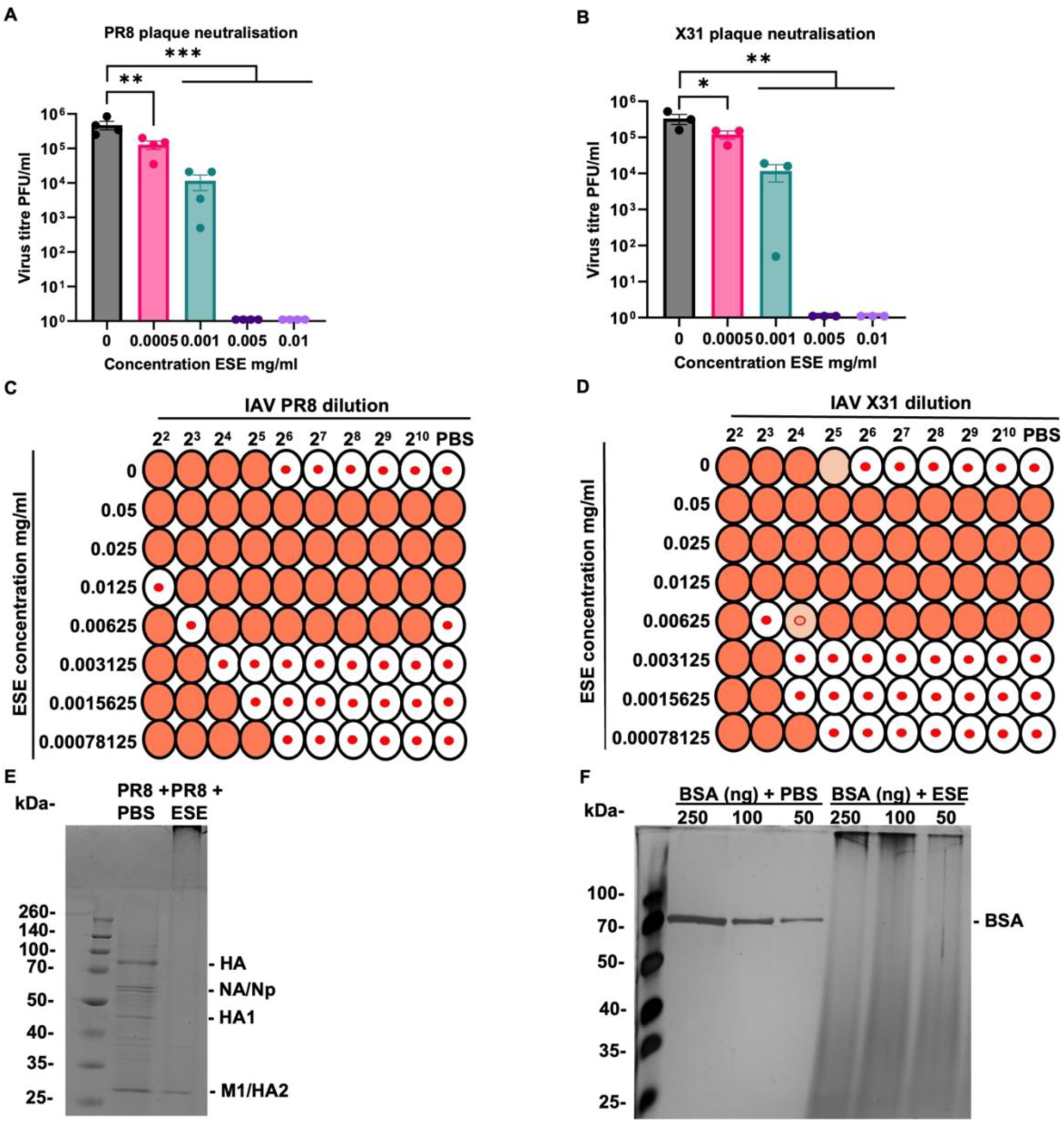
ESE interacts with hemagglutinin and causes protein aggregation. **(A) (B)** Plaque neutralisation. PR8 **(A)** and X31**(B)** were diluted to 10^6^ PFU/mL and incubated 1:1 with shown concentrations of ESE or infection media for one hour. Serial ten-fold dilutions were then performed and used to infect MDCK monolayers for one hour prior to addition of overlay and fixing and staining 3 days later. Data represented as mean ± SEM of three or more independent experiments. Asterisks indicate statistical difference (one-way ANOVA with Dunnett’s multiple comparisons test; *P<0.05, **P<0.01, ***P<0.001). **(C) (D)** HA assay. PR8 and X31 were serially diluted two-fold in PBS. ESE was also serially diluted two-fold and incubated in equal volumes with IAV for 30 minutes prior to addition of 0.5% chicken RBC. HA titres were then calculated. **(E) (F)** ESE causes aggregation of proteins. PR8 **(E)** or shown concentrations of BSA **(F)** were incubated with PBS or 2.5 mg/mL ESE prior to separation on an SDS-PAGE gel and Coomassie **(E)** or silver **(F)** staining.

### Virus cell binding is inhibited by ESE

The data so far suggests that ESE is binding to IAV, potentially preventing cell entry. This was further explored using transmission electron microscopy (TEM). Purified IAV PR8 was examined by TEM in the presence of ESE or PBS, using the negative staining technique. Large numbers of regularly shaped, 70 nm sized virions were detected in the control (PBS) sample. In the presence of high concentrations of ESE, virions were smaller, approximately 40 nm, and partly appeared fragmented (Figure 3A). This interaction is likely to inhibit cell binding and this was further explored using IAV PR8 labelled with the fluorescent dye Alexa fluor 488 (PR8-488). MDCK cells were chilled to 4°C prior to infection with PR8-488 pre-treated with ESE or infection media to allow virus binding but not internalisation. FACS analysis of cells shows pre-treatment of PR8-488 with 0.01 mg/mL ESE caused a significant reduction (79.3% mean reduction) in virus cell binding at 0.01 mg/mL (Figure 3C).

**Figure 3.**
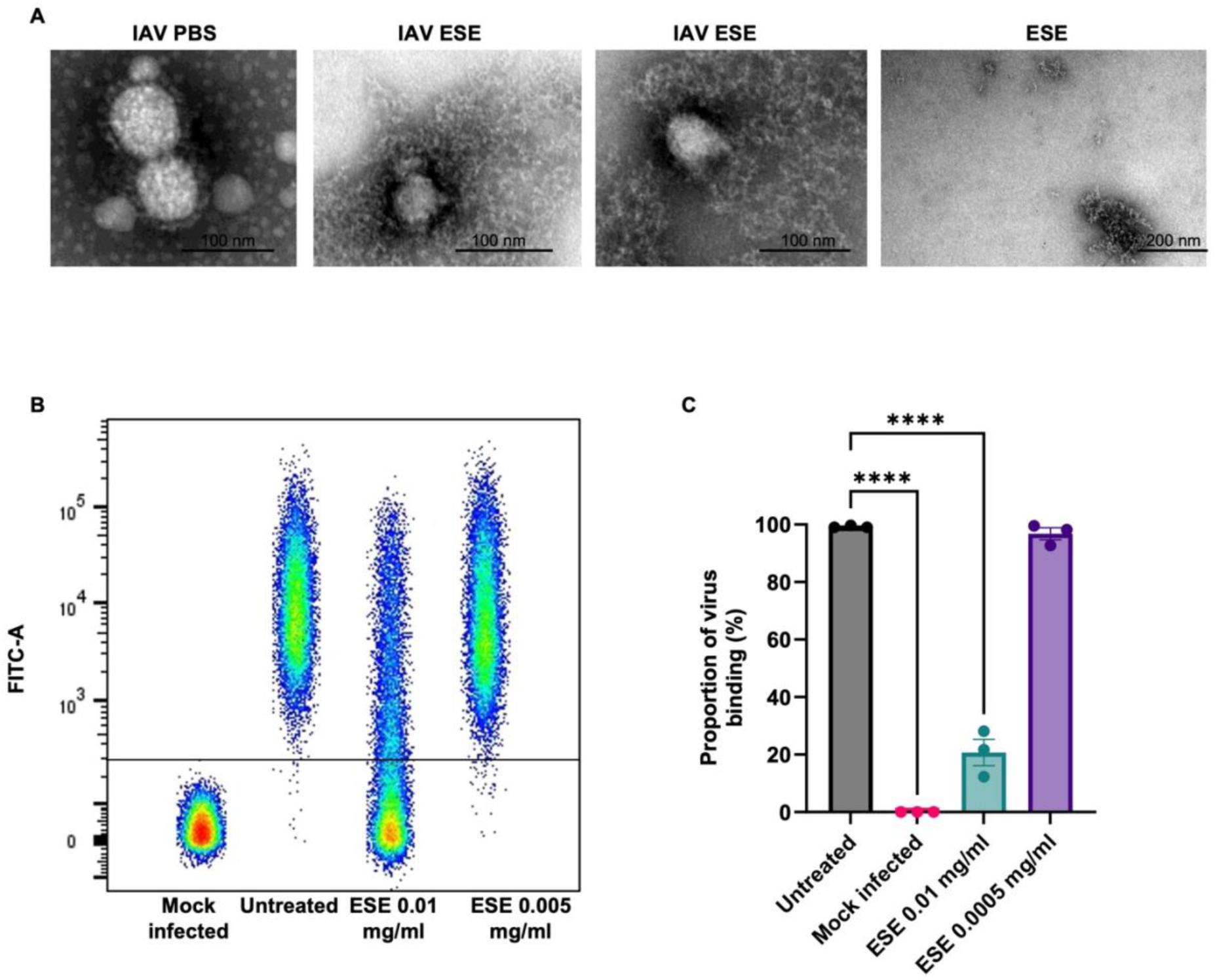
ESE interacts with IAV and prevents cell binding. **(A)** IAV PR8 in the presence of PBS or ESE, transmission electron microscopy, negative staining technique. IAV was purified through a sucrose cushion and mixed with PBS or 2.5 mg/mL ESE final and fixed with PFA. Left: IAV with PBS, showing an aggregate of regularly shaped virions of appr. 70 nm diameter. Middle: IAV with ESE. Virions are small (appr. 40 nm diameter) and appear partly fragmented. Right: ESE, without virions. **(B) (C)** FACS analysis of PR8-488 binding to MDCK cells. PR8-488 MOI 3 was incubated with infection media or shown concentrations of ESE for one hour prior to infection of MDCK cells for one hour at 4 degrees. Unbound virus was removed by PBS washing prior to fixation and analysis. **(B)** Dot plot representation of virus cell binding. **(C)** Proportion of virus binding calculated by FACS analysis. Data represented as mean ± SEM of three independent experiments. Asterisks indicate statistical difference (one-way ANOVA with Dunnett’s multiple comparisons test; ****P<0.0001).

### ESE promotes cell survival and reduces virus titre after virus cell entry

The initial screen of anti-influenza activity elucidated another anti-viral mechanism in events following virus cell binding. This was investigated using the same three strains of IAV and the addition of ESE one hour after infection, following cell entry. Cells were infected at a low MOI 0.001 to allow for multiple rounds of infection. Viable cells were measured by MTS assay at 72 hpi and given as a percentage of a mock infected control. ESE reduced virus induced cell death in a dose dependent manner in all three strains of IAV and no cytotoxicity was observed. To assess whether the promotion of cell survival was due to a reduction in virus titre, growth curves were performed. 0.01 mg/mL ESE reduced virus titre in all three strains, while 0.005 mg/mL also resulted in a reduction in virus titre. Interestingly, PR8 titre following addition of ESE appeared to increase over the 72 hours and was a similar titre to the untreated control at 72 hours which was not seen for X31 or Eng195 strains.

### Virus release and Neuraminidase activity is inhibited by ESE

Time of addition assay over one virus replication cycle (12 h) was carried out to assess which stage of the virus lifecycle was being affected by ESE. ESE appeared particularly effective at reducing IAV X31 virus titre and virus induced cell death (Figure 4C and 4D). Therefore, MDCK cells were infected with X31 MOI 0.01 on ice to allow virus binding but not internalisation. Virus entry was synchronised by rapidly increasing the temperature to 37 degrees and 0.01 mg/mL ESE was added at different time points (0, 2, 4, 6, 8, 10 hpi) (Figure 5A). Virus supernatant was collected at 12 hpi and cells lysed in Trizol following the completion of one virus lifecycle (12 hpi). Intracellular virus titres (Figure 5B), calculated by measuring viral load in cell lysate at 12 hpi by qRT-PCR, were compared to extracellular virus titres in the supernatant at 12 hpi titrated by plaque assay (Figure 5C). Viral load at 12 hpi was reduced when ESE was added at 0 hours following cell binding, although not significantly, while there was no noticeable decrease in viral load at 12 hpi when ESE was added at the remaining time points (Figure 5B). Interestingly, despite the high viral load, the release of infectious virus in the supernatant at 12 hpi was inhibited with the addition of ESE at 0, 2 and 4 hours while the titre was reduced with the addition of ESE at 6, 8 and 10 hours (Figure 5C). Influenza viruses utilise NA activity to cleave sialic acid at the cell surface from progeny virus, facilitating virus release. Therefore, the ability of ESE to inhibit NA activity of PR8, X31 and Eng195 was explored using the fluorescent neuraminidase substrate MUNANA with zanamivir, a known NA inhibitor, used as a positive control. ESE inhibited NA activity of all three strains of IAV, most effectively against X31 IC50 0.0005 mg/mL, followed by Eng195 IC50 0.0032 mg/mL and least effectively against PR8 IC50 0.0128 mg/mL. These values may explain the increased virus titre of PR8 at 72 hpi compared to X31 and Eng195 (Figure 4B).

**Figure 4.**
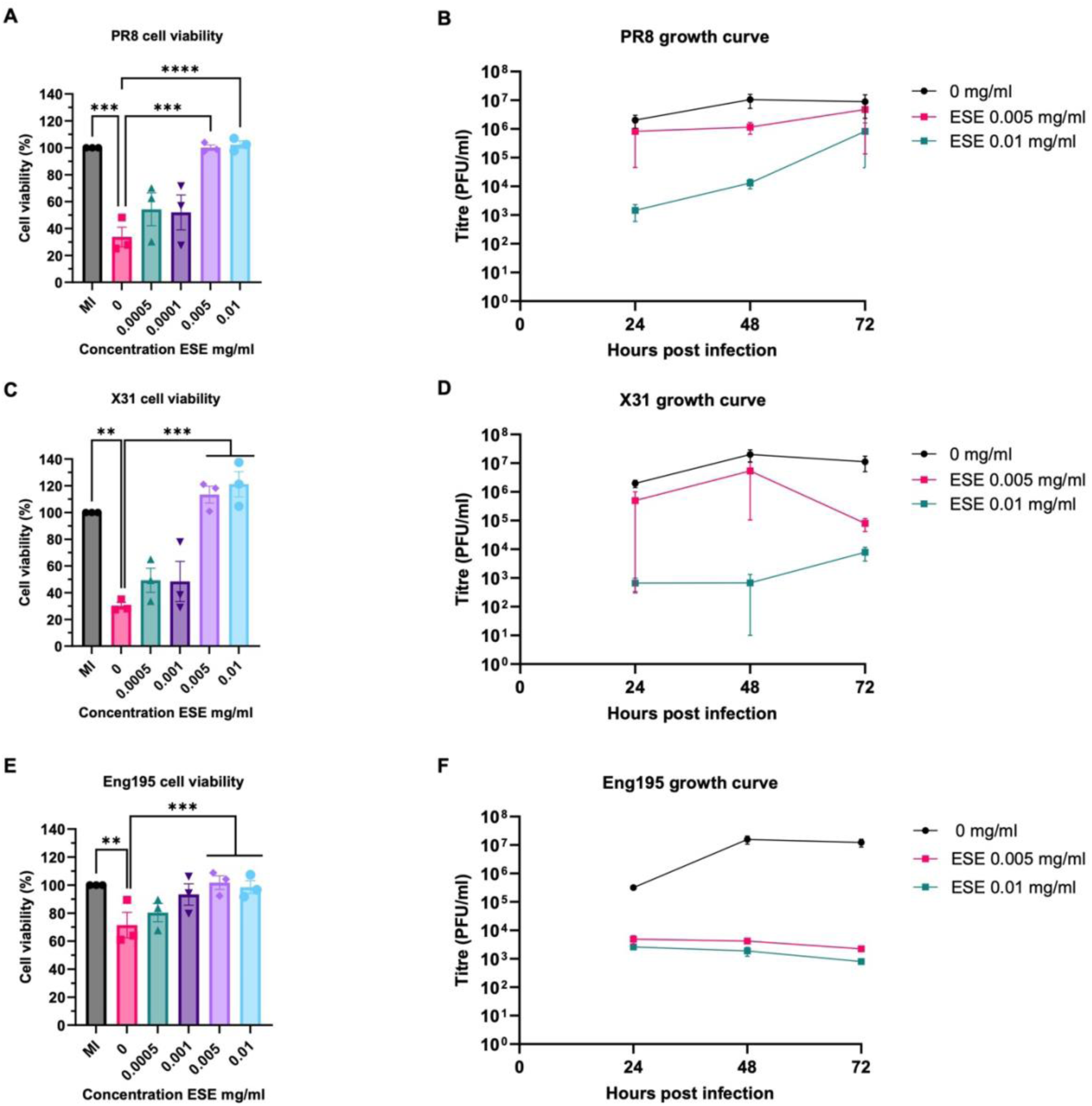
ESE promotes cell survival and reduces virus titre when added after infection. **(A) (C) (E)** MDCK cells were mock infected or infected with IAV PR8 (A), X31 (C) or Eng195 (E) MOI 0.001 for one hour. Inoculum was then removed and ESE concentrations or infection media were added. Cell viability was assessed by MTS assay 72 hours post infection and given as a percentage of a mock infected control (n=3). Data represented as mean ± SEM. Asterisks indicate statistical difference (two- way ANOVA with Dunnett’s multiple comparisons test; **P<0.01, ***P<0.001 ****P<0.0001). **(B) (D) (F)** IAV growth curves. MDCK cells were infected with IAV PR8 (B), X31 (D) or Eng195 (F) MOI 0.001 for one hour. Inoculum was then removed and ESE concentrations or infection media were added. Supernatant was harvested at indicated time points and titrated by plaque assay on MDCK cell monolayers. Data represent the mean value ± SEM (n=3).

**Figure 5.**
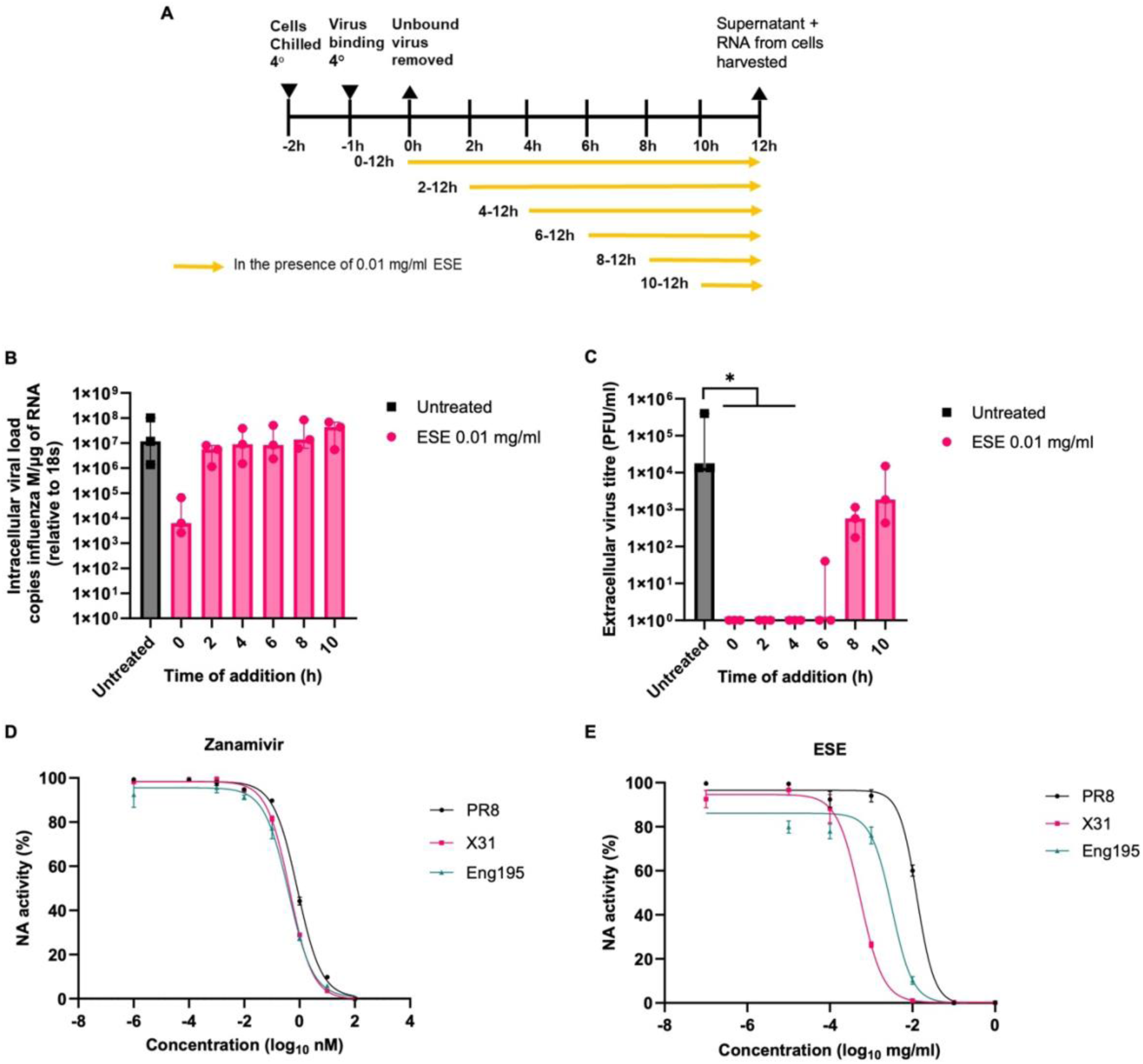
IAV release is inhibited by ESE. **(A)** Schematic of time of addition assay. MDCK cells were chilled to 4 degrees and infected IAV X31 MOI 0.01 for one hour on ice to allow virus binding. Temperature was then increased to 37 degrees to allow synchronised entry and 0.01 mg/mL ESE was added at indicated time points. At 12 hours post infection, cells were lysed with trizol and supernatant collected. Viral load in cells **(B)** and titre in supernatant **(C)** were measured by qRT-PCR and plaque assay respectively). Data represent the median value + interquartile range (IQR) of three independent experiments. Asterisks indicate statistical difference (Kruskal-Wallis with Dunn’s multiple comparisons test; *P<0.05). **(D) (E)** Fluorescent neuraminidase substrate (MUNANA) based neuraminidase inhibition assay using Zanamivir **(D)** or ESE **(E)** (n=3).

### Nuclear import and internalisation of IAV nucleoprotein is reduced

The reduction in viral load when ESE was added post binding indicated a disruption of the virus lifecycle prior to replication. The localisation and internalisation of input IAV nucleoprotein (NP) was therefore explored in the presence of a protein synthesis inhibitor, cycloheximide. Briefly, MDCK cells were chilled to 4 ℃ to allow virus binding but not entry. IAV X31 was added at MOI 3 and allowed to bind for one hour. ESE or the endosome acidification inhibitor ammonium chloride (NH_4_) were then added, and virus entry synchronised at 37 ℃ for 3 hours. Cells were then fixed and stained for IAV nucleoprotein (NP). In the untreated sample, NP localised to the cell nucleus as expected while NH_4_ prevented localisation of NP to the cell nucleus and reduced the number of puncta (Figure 6A). ESE also appeared to reduce internalisation with a decrease in puncta detected while localisation of NP to the cell nucleus also appeared to be reduced compared to the untreated control (Figure 6B and 6C).

**Figure 6.**
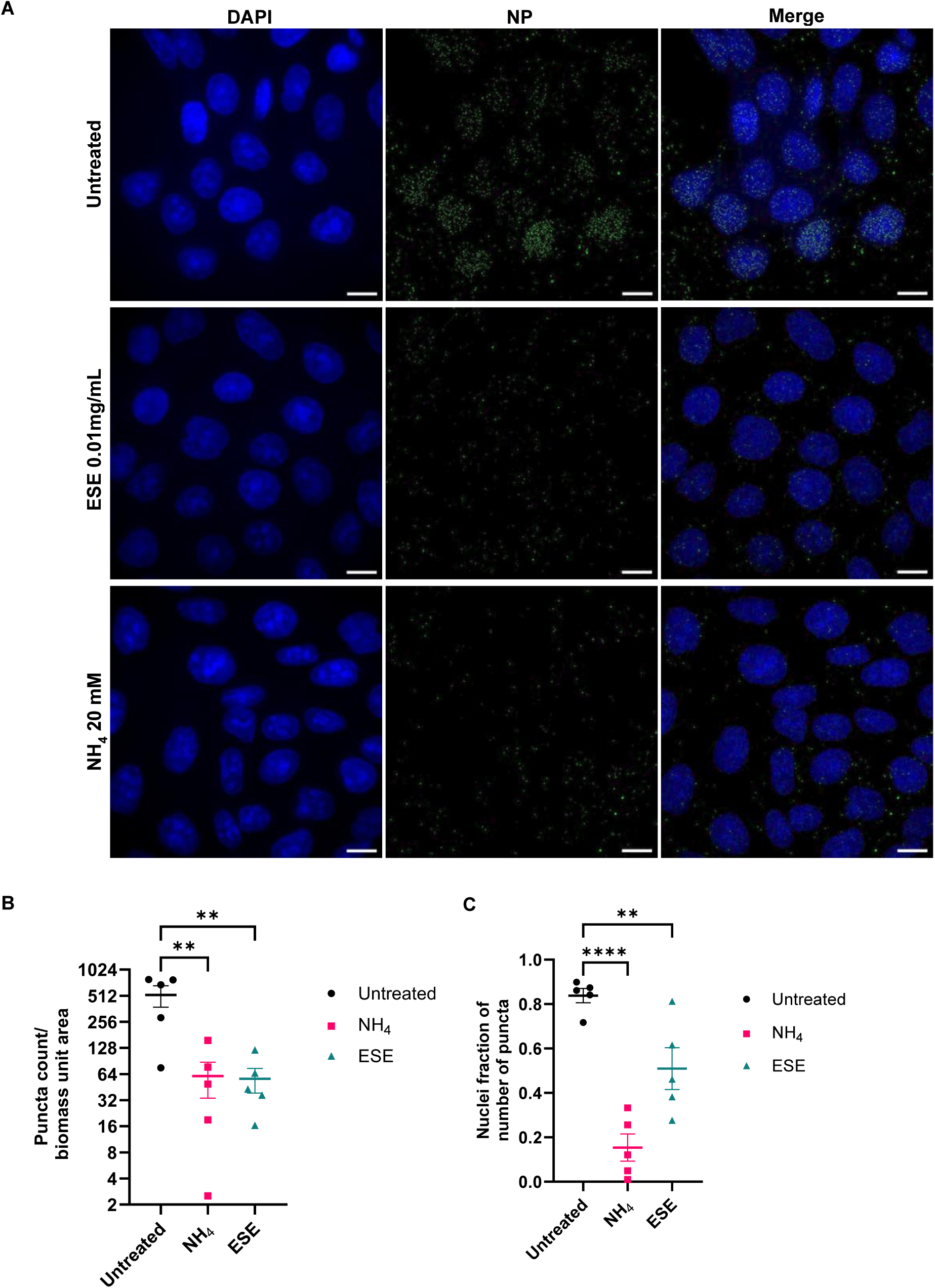
ESE reduces internalisation of IAV NP. **(A)** MDCK cells were grown to 80% confluency and chilled at 4 degrees for 90 minutes. X31 at an MOI of 3 was added to cells for one hour at 4 degrees in the presence of 1 mM cycloheximide. Unbound bound virus was then removed with 3 washes of ice-cold PBS prior to the addition of infection media, 20mM NH_4_ or 0.01 mg/mL ESE. Cells were rapidly warmed to 37°C to allow synchronised entry in 5% CO_2_ for 3 hours prior to fixing and permeabilization with 100% methanol. IAV NP was detected with a mouse anti-NP and an Alexa Fluor 488-conjugated goat anti-mouse secondary antibody. Bars = 10 μM **(B)** Puncta count and nuclei fraction **(C)** were calculated from 5 fields of view chosen at random representing over 100 cells. Data represented as mean ± SEM. Asterisks indicate statistical difference (one-way ANOVA with Dunnett’s multiple comparisons test; **P<0.01, ****P<0.0001)

### Investigating ESE effectiveness on 3-D model

The effectiveness of ESE to inhibit IAV infection of human bronchial epithelial cells (HBEC3 - KT) grown at the air liquid interface was explored. Cells were allowed to differentiate for 14 to 21 days prior to infection and differentiation was confirmed by endpoint PCR of differentiation markers MUC5AC, CBE1, SCGB1A1, BPIFA1 and TEKT1 (data not shown). Inhibition of virus cell entry was investigated by infecting cells with 10^5^ PFU IAV Eng195 in the presence of ESE or PBS at the apical surface and virus titre calculated from apical washes at 24 and 48 hours. A significant reduction in virus titre was observed at both 24 and 48 hpi compared to the PBS treated control (Figure 7A and 7B). The potential of ESE to inhibit infection following virus entry was investigated next, with cells again being infected with 10^5^ PFU IAV Eng195 and ESE or PBS being dosed repeatedly at the apical surface at 2 hpi, 6 hpi, 24 hpi and 30 hpi. Virus titre was again calculated from apical washes at 24 and 48 hpi. A significant reduction in virus titre was observed at both 24 and 48 hpi compared to the PBS treated control (Figure 7C and 7D).

**Figure 7.**
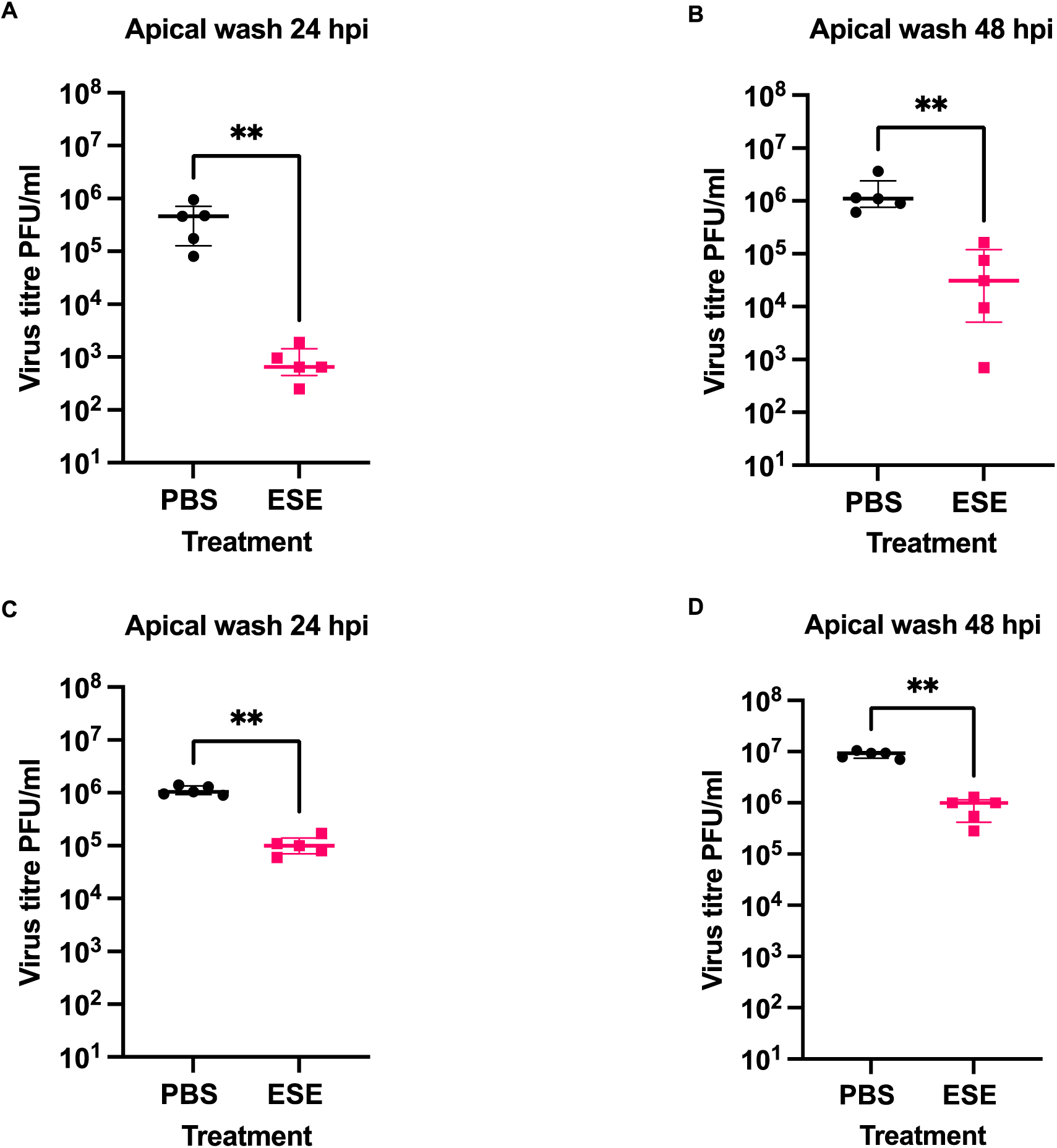
Investigating ESE effectiveness on a differentiated 3-D model. **(A**) **(B)** HBEC3-KT cells were grown at the air liquid interface for 14 to 21 days. Differentiation was confirmed by endpoint PCR. Cells were infected at the apical surface with 10^5^ PFU IAV ENG195 in the presence of 0.01 mg/mL ESE or PBS. Apical washes were performed at 24 **(A)** and 48 **(B)** hours and virus titre calculated by plaque assay **(C) (D)** Differentiated HBEC3-KT cells were infected at the apical surface with 10^5^ PFU IAV ENG195. 0.01 mg/mL ESE or PBS was added to the apical surface at 2 hpi, 6 hpi, 24 hpi and 30 hpi. Apical washes were performed at 24 **(C)** and 48 **(D)** hours and virus titre calculated by plaque assay. Data represent the median value + IQR of 5 biological replicates. Side-by-side comparisons were made using Mann-Whitney U test (** represents p<0.005).

### Intranasal administration of ESE reduces viral load in mice infected with IAV

The effectiveness of ESE intranasal administration was examined in a mouse model. In a first step, we determined whether ESE had any pathological effect when administered intranasally. For this, 6-8 week old female C57Bl/6J mice were treated intranasally with 5 mg/kg ESE, either as one dose in 50 µl PBS to ensure that a sufficient amount of ESE reaches the lung parenchyma, i.e., the alveoli, or as 5 doses in 10 µl PBS on 5 consecutive days. All mice were euthanised at day 4 post adminstration. Mice receiving ESE in one dose in 50 μl PBS exhibited mild weight loss at day 1 (Supplementary Figure S2) but then showed progressive weight gain. In contrast, mice receiving lower daily doses in 10 μl basically maintained their weights throughout the course of the experiment (Supplementary Figure S2). The post mortem examination did not reveal any gross pathological changes in the animals. All relevant organs/tissues were subjected to a histological examination. This did not reveal any pathological changes in any organs apart from the lungs of the mice that had received ESE in one 50 µl dose. In all four mice, the lungs exhibited mild inflammatory changes, represented by focal granulomatous (i.e. macrophage dominated) infiltrates; this likely represents a response to inhaled ESE (Supplementary Figure S3). These findings indicate that intranasal ESE application has no systemic pathological effect but show that ESE can induce a mild foreign body reaction in the lung when applied intranasally in a larger fluid volume. This is not seen when most of the ESE can be expected to remain in the upper respiratory tract, i.e. when applied in lower volumes. However, the inflammatory reaction was only mild and did not induce any clinical signs as all mice had gained weight by two days after instillation which can be considered the shortest time span for a granulomatous reaction to have developed, since uptake of suitable inhaled material by alveolar macrophages alone takes 6-12 hours [24]. The results of the histological examination of the individual animals are provided in Supplementary Table S2.

For the subsequent infection experiment, ESE was applied at the above-mentioned dose, and in 10 µl PBS, to avoid any ESE induced inflammatory reaction. Briefly, mice were infected intranasally with a sub lethal dose of IAV X31. Three ESE treatment approaches were taken, prophylactical (treatment start at 2 h pre infection), at the time of infection, and therapeutical (treatment starting at 3 hpi); treatment then continued daily until sacrifice at day 5 (Figure 8A). Prophylactical cohort and time of infection cohort were also dosed again at 3 hpi. Control mice received daily doses of PBS starting at 3hpi.

**Figure 8.**
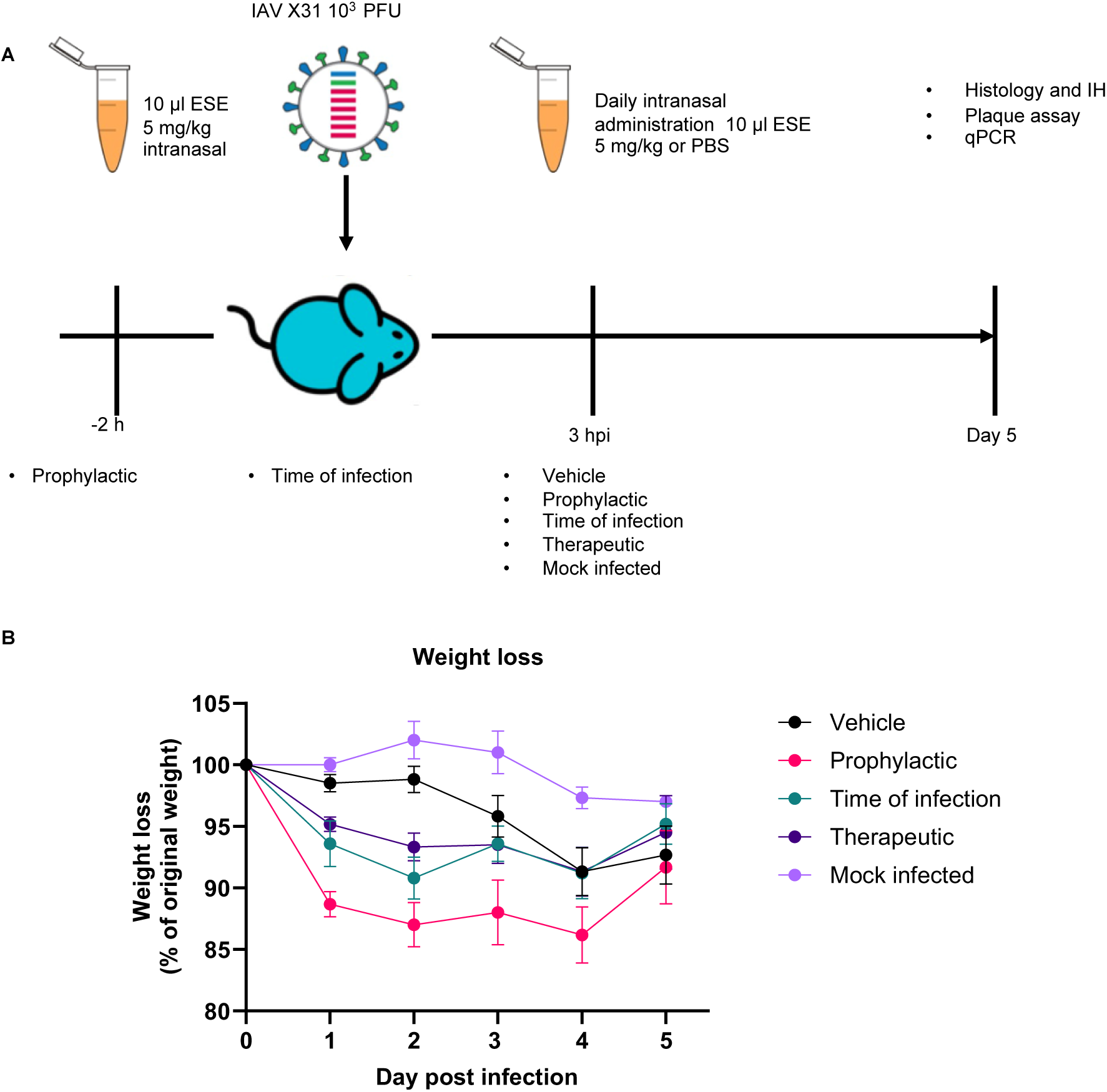
Intranasal administration of ESE *in vivo*. **(A)** Schematic of *in vivo* infection and treatment administration. Female C57BL/6 mice were challenged intranasally with 10^3^ PFU IAV X31 in 10µl. Daily intranasal administration in 10µl of 5mg/kg ESE or PBS started at indicated time points and continued until day 5 where mice were sacrificed by cervical dislocation. **(B)**. Mice were monitored for weight loss at indicated time points (n = 6). Data represent the mean value ± SEM. Comparisons were made using a repeated measures two-way ANOVA (Supplementary table S1).

Vehicle treated infected animals showed the typical weight change after IAV infection, with obvious weight loss by day 3 and the peak at day 4 [25]. With ESE treatment, a drop in weight was observed at day 1 prior to virus mediated eight loss which starts at day 3; this was most intense in the prophylactic group where it remained rather stable at the overall lowest level until day 4 (Figure 8B). This initial drop in weight might be the consequence of the repeated anesthesia (3 times) this group of mice was subjected to on day 0 as this weight loss was not seen previously when administering ESE (Supplementary Figure S2). The trend was similar in the other two treatment groups, and at day 5 the weights of the various groups did not differ significantly (Supplementary Table S1).

Examination of the animals at the end of the experiment, i.e. day 5 post infection, confirmed IAV infection in all mice. They all harboured viral RNA in the nasal tissue, as determined by qRT-PCR for the viral M gene (Figure 9A). Interestingly, the viral load was significantly higher in both the therapeutic (median 1.83 x 10^7^ copies of M/μg RNA) and prophylactic (median 3.43 x 10^7^ copies of M/μg RNA) treatment cohorts compared to the vehicle only cohort (median 5.60 x 10^6^ copies of M/μg RNA), whereas it was significantly lower in mice for which the treatment had started at the time of infection (median 4.04 x 10^3^ copies of M/μg RNA). The latter group also exhibited the lowest viral loads (median 222 copies of M/μg RNA) in the lungs (Figure 9B) from which infectious virus could not be isolated when the virus titre was determined by plaque assay performed on homogenised lung tissue (Figure 9C). Significantly lower viral loads (median 5.05 x 10^5^ copies of M/μg RNA) as well as significantly lower virus titres (median 1.52 x 10^4^ PFU/lung) were also found in the lungs of mice that had received the therapeutic treatment compared to vehicle (median 1.64 x 10^7^ copies of M/μg RNA and 4.70 x 10^5^ PFU/lung) controls (Figure 9B and 9C). The prophylactic treatment regime resulted in a reduction in viral loads (median 3.46 x 10^6^ copies of M/μg RNA) and in the virus titre in the lungs (median 6.13 x 10^4^ PFU/lung), although the difference to mice receiving vehicle did not reach significance (P=0.0584). It has been previously reported that seaweed derived compounds can increase the interferon response to infection [18]. This was explored by analysing ISG15 expression. ISG15 expression in lungs (Figure 9D) was significantly lower in all cohorts compared to the vehicle only cohort and is supportive of decreased infection in mice intranasally treated with ESE.

**Figure 9.**
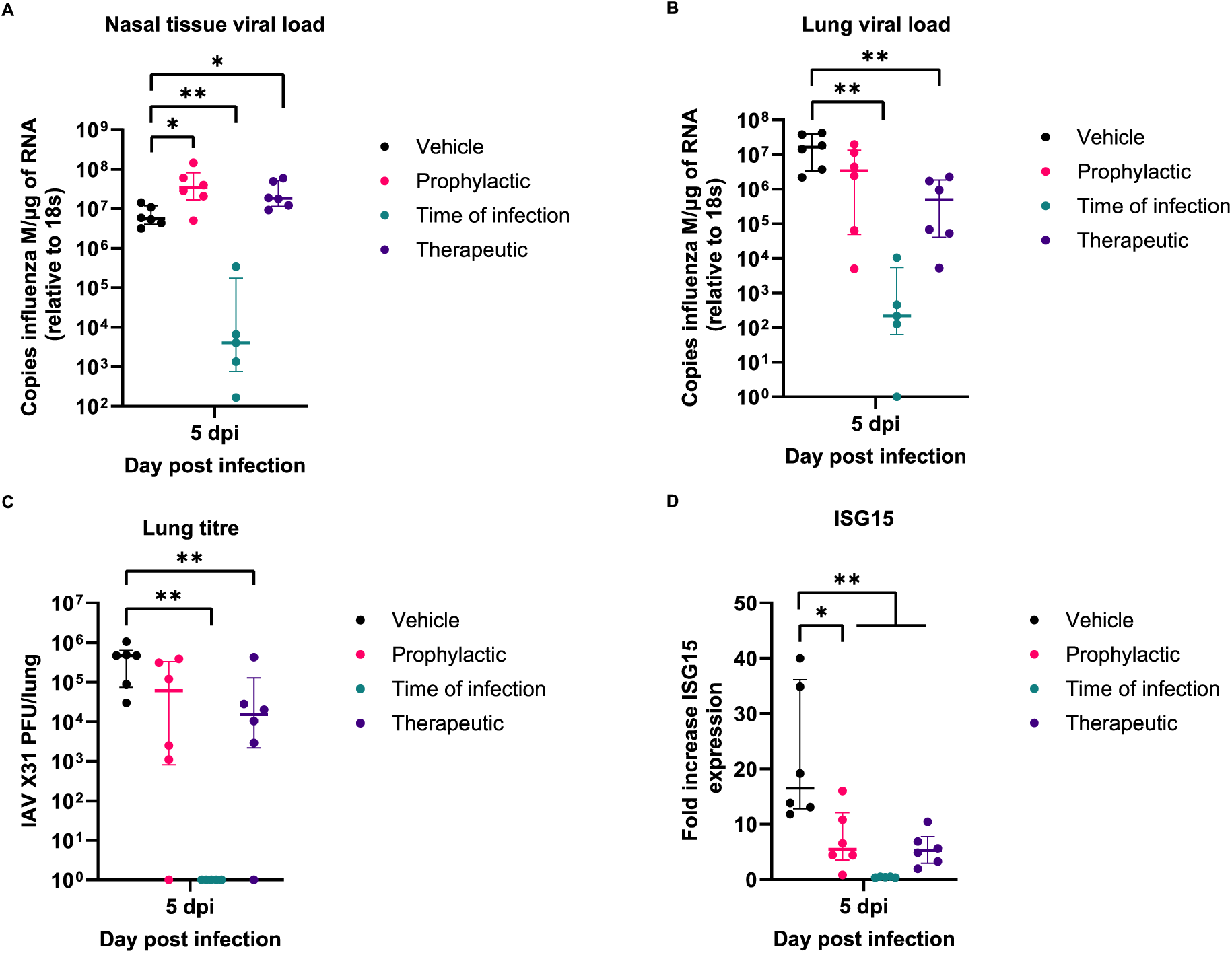
Viral load in mice challenged with IAV. Viral load in RNA extracted from nasal tissue **(A)** or lungs **(B)** were calculated by qRT-PCR of Influenza M gene and normalised relative to levels of 18S rRNA (n=6). **(C)** IAV titre from homogenised right lung lobe was calculated by titration on MDCK cell monolayers (n=6). **(D)** ISG15 gene expression was calculated from by qRT-PCR of lung RNA using delta delta ct method and given as a fold increase of mock infected mice (n=6). Data represent the median value + IQR. Side-by-side comparisons were made using Mann-Whitney U test (* represents p < 0.05, ** represents p<0.005).

The histological and immunohistological examinations of the vehicle treated infected mice yielded the typical changes elicited by IAV X31 at this time point [25], [26] representing a necrotic bronchitis/bronchiolitis and a multifocal acute desquamative pneumonia, with virus infection of both respiratory and alveolar epithelial cells, and virus antigen in macrophages (Figures 10A and 11A). Both the prophylactic and therapeutic treatment cohorts in the majority exhibited the same changes, though less extensively (Figure 10B and 10D), with evidence of free virus admixed with fluid in the lumen of bronchioles (Figure 11B and 11C). In contrast, virus antigen was not detected in the lung of the animals subjected to the time of infection treatment regime. Most lungs showed no or only minimal inflammatory changes (Figure 10C); however, in one animal (#3.3), there was evidence of focal hyperplasia of the bronchiolar epithelium and a moderate (pyo)granulomatous pneumonia, suggesting that the lung had been infected. Overall, the findings suggest that pulmonary infection is blocked when ESE is applied at the time of infection. Detailed information on individual animals is provided in Supplemental Table S3.

**Figure 10.**
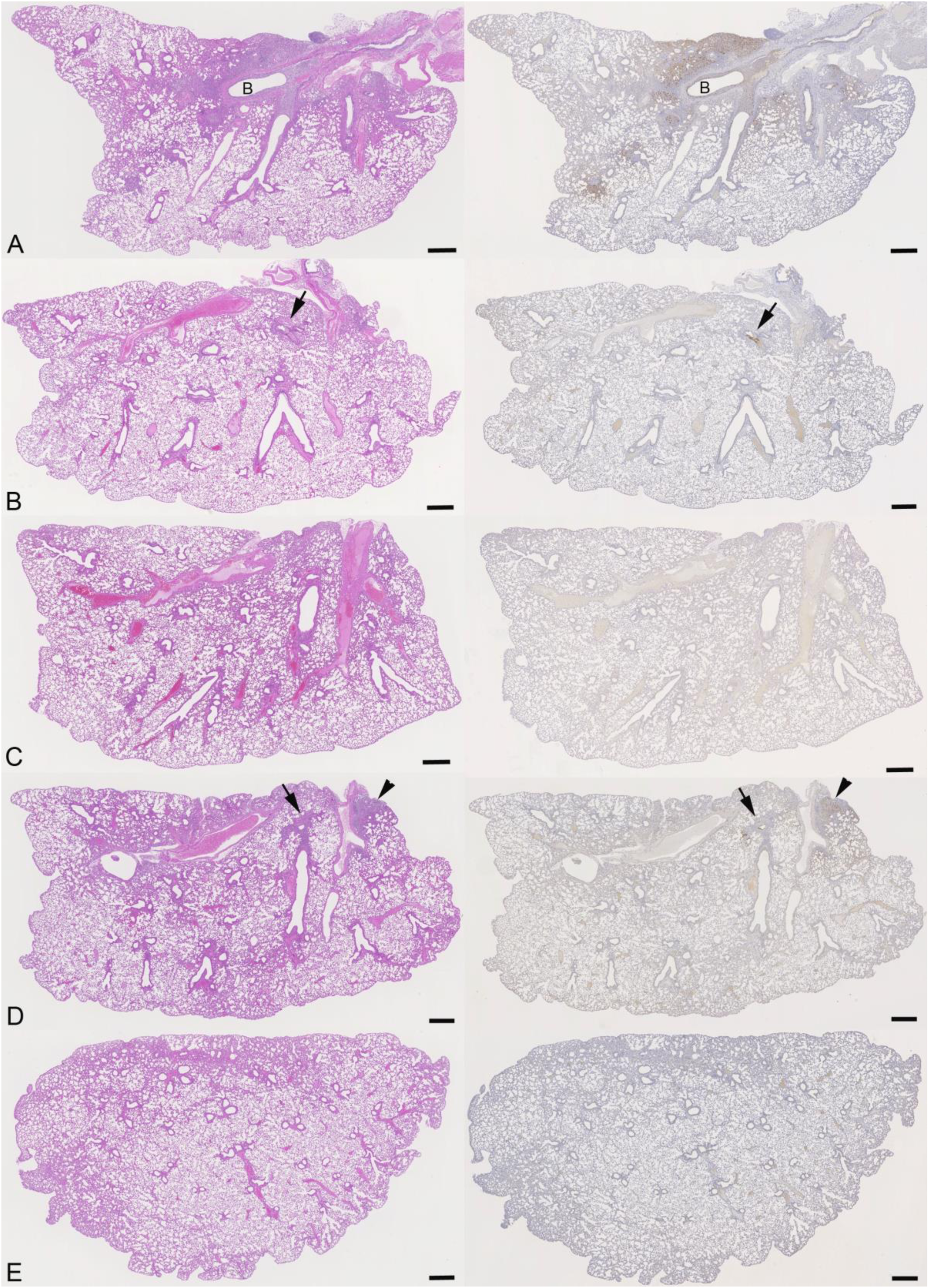
Histological features and viral antigen expression in mice challenged with IAV and treated with ESE. **(A)** Animal treated with PBS (#1.6). Necrotic bronchitis and bronchiolitis with extensive viral antigen expression and desquamative pneumonia in adjacent parenchymal areas (for higher magnification, see Figure 12A). B: bronchus. **(B)** Animal of prophylactic treatment scheme (#2.6). Changes are restricted to one bronchiole (arrow) with a patch of infected, IAV antigen positive epithelial cells and some material in the lumen (for higher magnification, see Figure 12B). **(C)** Animal of time of infection treatment scheme (#3.6). The lung parenchyma appears unaltered, and there is no evidence of viral antigen expression. **(D)** Animal of therapeutic treatment scheme (#4.6). Focal area with peribronchial leukocyte infiltration and small bronchioles with infected epithelial cells (arrow), and focal parenchymal area with changes consistent with desquamative pneumonia and viral antigen expression in numerous alveolar epithelial cells and macrophages (arrowhead). **(E)** Mock infected control animal treated with PBS (#5.2). The lung parenchyma is unaltered, and there is no evidence of viral antigen expression. Left column: HE stain; right column: immunohistology, hematoxylin counterstain. Bars = 500µm.

**Figure 11.**
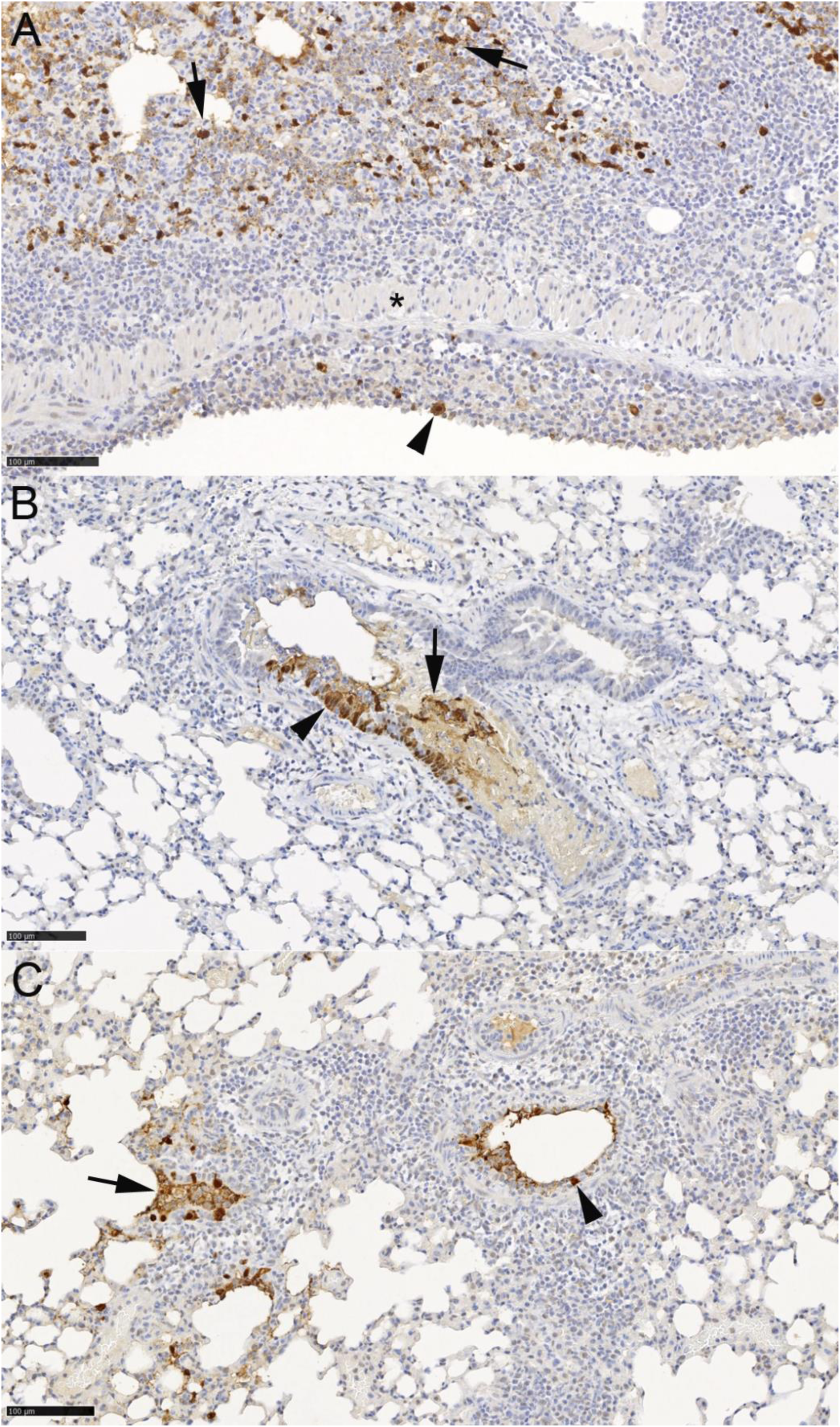
Viral antigen expression in the lungs of mice challenged with IAV, treated with ESE and examined at day 5 post infection. Detailed individual animal data is provided in Supplementary Table S3. **(A)** Animal treated with PBS (#1.6). Bronchiole with necrosis of epithelial cells and IAV antigen expression in sloughed off degenerate cells (arrowhead). Extensive viral antigen expression in alveolar epithelial cells and macrophages (arrows) in an adjacent parenchymal area with changes consistent with desquamative pneumonia. **(B)** Animal of prophylactic treatment scheme (#2.6). Bronchiole with patch of infected, IAV antigen positive epithelial cells (arrowhead) and positive material in the lumen (arrow). **(C)** Animal of therapeutic treatment scheme (#4.5). Small bronchioles with intact (arrowhead) and degenerate infected epithelial cells and some positive material in the lumen. Immunohistology, hematoxylin counterstain. Bars = 100µm.

## Discussion

This study focused on a seaweed extract isolated from the seaweed *Ascophyllum nodosum* and its potential use as a novel anti-viral agent derived from a renewable source. ESE inhibited the three strains of IAV tested most effectively when combined with virus first. This interaction was not reversed by serial dilution and appears to cause protein aggregation. In contrast, ESE pre-treatment of MDCKs did not reduce virus titres suggesting ESE may not interact with cells directly. The ultrastructural examination provided further evidence of this interaction between ESE and IAV while FACS analysis using fluorescently labelled virus deduced ESE inhibits infection by blocking virus cell binding and therefore entry.

Interestingly, ESE appears to have more than one mode of action. The addition of ESE to cells following virus cell entry reduced viral titres and protected cells from virus induced cell death. Time of addition assay over one replication cycle indicated that replication was not being inhibited but rather release of infectious virus. This was confirmed by the ability of ESE to inhibit NA activity in the three IAV strains, activity of which is utilised by IAV to facilitate progeny virus release from the cell surface through cleavage of sialic acid [10], [11]. Furthermore, ESE appears to reduce virus internalisation following cell binding.

The therapeutic potential of ESE was explored utilising human 3-D airway models grown at the air liquid interface. These displayed reduced virus titres when ESE was added to the apical surface before or following infection. Using a murine model of IAV infection, we could show that intranasally applied ESE is well tolerated when applied in a low amount of fluid rather than being instilled in a larger volume that immediately reaches the alveoli and induces a mild granulomatous reaction. Administration of ESE in mice starting at the time of infection or 3 hours after infection caused a significant reduction in viral load in the lungs. It was lowest in the former cohort from which infectious virus could not be isolated from the lungs at all. Interestingly, viral load appeared higher in the nasal tissue for the therapeutic and prophylactically treated cohorts, suggesting ESE could be forming a protective barrier in the nasal tissue reducing the amount of virus reaching the lungs, indicating a potential for nasal application of ESE.

To conclude, ESE displays anti-IAV activity *in vivo* and *in vitro*, preventing virus cell binding and inhibiting release of progeny virus by targeting viral neuraminidase activity. Given the nature of inhibition of virus cell binding, it is possible ESE will display antiviral efficacy against other respiratory viruses and should be investigated further. ESE displays broad-spectrum anti-IAV activity and may have potential to be developed into a nasal spray for the prophylactic or therapeutic treatment of influenza infections.

## Methods

### Seaweed extract

Enriched Seaweed Extract (ESE) was produced by hydroethanolic extraction of fresh *Ascophyllum nodosum* followed by enrichment using C_18_ solid phase extraction. This afforded a phlorotannin rich extract as previously described [22]. ESE was dissolved in phosphate buffered saline (PBS) to a final concentration of 10 mg/mL. This stock solution was sterile filtered through a 0.2 μM filter and stored at 4 °C prior to use.

### Virus, cell lines and media

Madin Darby Canine kidney (MDCK) and MDCK-SIAT1 cells were maintained in DMEM supplemented with 10% Foetal bovine serum (FBS) at 37 °C in 5% CO_2_. MDCK-SIAT1 cells were further supplemented with 1 mg/mL geneticin (G418). Influenza A Virus A/Puerto Rico/8/1934 H1N1 (PR8), Influenza A Virus A/X-31 H3N2 (X31) and Influenza A Virus A/England/195/2009 H1N1 (Eng195) were propagated on MDCK or MDCK-SIAT1 cells.

### Plaque reduction assays

IAV strains (PR8, X31, Eng195) were diluted to 25-50 PFU. For the cell-pre-treatment assays, cells were treated with ESE diluted in serum free DMEM supplemented with TPCK-trypsin (infection media) for one hour. ESE was removed and cells washed with PBS. Virus was added for one hour to allow for internalisation prior to addition of an overlay. Virus pre-treatment assays, virus was incubated 1:1 with ESE or infection media prior to infection of cell monolayers for one hour. Post entry events were investigated with the addition of ESE to plaque assay overlays 1 hour post infection of cell monolayers. Plaques were given as a percentage of an untreated control.

### Growth curves

Cells were grown to 80% confluency in 12 well plates and washed twice with PBS prior to infection MOI 0.001 for one hour at 37 °C in 5% CO_2_. Virus inoculum was removed and infection media or infection media containing ESE was added. Virus supernatant was harvested at indicated timepoints and titrated by plaque assay on cell monolayers

### Cell viability assays

Cells were grown to 80% confluency in 96 well plates and infected MOI 0.001 for one hour. Virus inoculum was removed and infection media containing ESE was added. Cell viability was measured by MTS assay (Promega) 72 hours post infection and given as a percentage of the untreated control.

### Time of addition assays

MDCK cells were chilled to 4 degrees for 90 minutes and infected with IAV X31 MOI 0.01 for one hour on ice to allow virus binding. Temperature was then increased to 37 degrees to allow synchronised entry and 0.01 mg/mL ESE was added at indicated time points. At 12 hours post infection supernatant was collected and cells lysed with TRIzol. Viral load in cells was measured by qRT-PCR of the IAV M gene and titre calculated by plaque assay.

### IAV nucleoprotein localisation

MDCK cells were grown to 80% confluency and chilled at 4 degrees for 90 minutes on coverslips. X31 at an MOI of 3 was added to cells for one hour prior at 4 degrees in the presence of 1 mM cycloheximide (Abcam). Unbound bound virus was then removed with 3 washes of ice-cold PBS prior to the addition of infection media, 20mM NH_4_ or 0.01 mg/mL ESE in the presence of 1 mM cycloheximide. Cells were rapidly warmed to 37°C to allow synchronised entry in 5% CO_2_ for 3 hours prior to fixing and permeabilization with 100% methanol. IAV NP was detected with a mouse anti-NP (Abcam) and an Alexa Fluor 488- conjugated goat anti-mouse secondary antibody. Coverslips were mounted onto slides using ProLong™ Gold Antifade Mountant with DAPI (Thermofisher) and imaged in lattice SIM mode using a Zeiss Elyra 7. Five fields of view were chosen at random consisting of over 100 cells. Images were SIM^2^ post-processed to generate super resolved and widefield images.

### Neuraminidase inhibition assay

Virus stocks were titred by a neuraminidase (NA) activity assay (Applied Biosystems, 4457091) based off NA activity of 5 μM 4-MU/60 minutes at 37 degrees. NA inhibition was calculated using a MUNANA-based neuraminidase assay (Applied Biosystems, 4457091). ESE or Zanamivir (Sigma) were serially diluted and incubated with virus for 30 minutes prior to addition of NA-Fluor substrate for 60 minutes. The plate was read using an excitation wavelength range of 350 nm to 365 nm and an emission wavelength of 440 nm to 460 nm. IC_50_ values were calculated using sigmoidal curve-fitting.

### Animal work

Animal work was reviewed and approved by the local University of Liverpool Animal Welfare Committee and performed under UK Home Office Project Licences PP4715265. Mice were all specified pathogen-free and maintained under barrier conditions in individually ventilated cages. Six to eight week old female C57BL/6J mice were purchased from Charles River (Margate, UK). Mice were randomly assigned into cohorts.

For the administration of ESE, mice were lightly anaesthetized with isoflurane and 5 mg/kg ESE in PBS was administered intranasally.

For the ESE tolerability study, two cohorts of 4 mice each were used; cohort 1 received one dose of ESE in 50 μl PBS at day 1 while cohort 2 received 5 doses of ESE in 10 μl PBS on 5 consecutive days; animals were euthanised at day 5 by cervical dislocation. They were dissected immediately after death and samples from all major organs and tissues collected and fixed in 10% buffered formalin.

For the infection study, 5 cohorts of 6 mice were used. vehicle or mock-infected mice received 10 µl of PBS. Administration of 5 mg/kg ESE in 10 μl PBS started at different time points depending on the cohort. For the prophylactic cohort, treatment started 2 h prior to infection, for the time of infection cohort, ESE was administered together with virus, and for the therapeutic cohort, treatment started 3 hpi. PBS treatment for the vehicle and mock-infected mice started 3 hpi. All prophylactic and time of infection cohort mice were treated again at 3 hpi, and all mice were treated daily thereafter.

For virus infection, mice were anesthetized lightly with KETASET i.m. and challenged intranasally with 10^3^ PFU IAV X31 in 10 μl sterile PBS or were mock-infected with the same volume of PBS. Mice were sacrificed at day 5 post infection. Lungs were removed immediately and the right lung snap frozen prior to downstream processing for virology. The left lung was fixed in 10% buffered formalin for 48 h. For histological and immunohistological examination, the lungs were routinely paraffin wax embedded.

### Lung homogenisation, RNA extraction and DNase treatment

The upper lobe and the lower lobe of the right lung were homogenised in 1ml TRIzol reagent or PBS for RNA extraction or titration by plaque assay respectively. Tissues were homogenised using stainless steel beads and a TissueLyser (Qiagen). Cell culture experiments were lysed in 0.5 mL of TRIzol reagent. The homogenates were clarified by centrifugation at 12,000xg for 5 min before full RNA extraction was carried out according to manufacturer’s instructions. RNA was quantified and quality assessed using a Nanodrop (Thermofisher) before DNase treatment using the TURBO DNA-free™ Kit (Thermofisher) as per manufacturer’s instructions.

### qRT-PCR for viral load

Viral loads were quantified as previously described [25] using the GoTaq® Probe 1-Step RT-qPCR System (Promega). The IAV primers and probe sequences are published as part of the CDC IAV detection kit using the following primers F: GACCAATCCTGTCACCTCTGAC, R: AGGGCATTTTGGACAAAGCGTCTA and Probe: 56-FAM CGTGCCCAGTGAGCAAGGACTGCA 3IABkFQ. The IAV reverse genetics plasmid encoding the M gene was used to serve as a standard curve. The thermal cycling conditions for all qRT-PCR reactions were as follows: 1 cycle of 45°C for 15 min and 1 cycle of 95°C followed by 40 cycles of 95°C for 15 sec and 60°C for 1 minute. Viral load was normalised relative to 18S rRNA. The 18S standard was generated by the amplification of a fragment of the murine 18S cDNA using the primers F: ACCTGGTTGATCCTGCCAGGTAGC and R: AGC CAT TCG CAG TTT TGT AC prior to purification using a QIAquick gel extraction kit (Qiagen). ISG15 gene expression was quantified using a SYBR Green-based real-time RT-PCR kit (Qiagen), ISG15 QuantiTect primers (Qiagen, QT00322749) and normalised to 18S rRNA using 18S QuantiTect primers (Qiagen, QT02448075).

### Infection of HBEC3-KT cells grown at air liquid interface

Immortalised human bronchial epithelial cells (HBEC3-KT) were grown on 12 mm transwells with 0.4 µm pore inserts (StemCell) in complete PneumaCult Ex Pluis media (StemCell) until confluent. Cells were then air lifted and basal media replaced with complete PneumaCult ALI medium (StemCell) for 14 to 21 days. Differentiation was confirmed by RT-PCR of differentiation markers. Differentiated cells were infected at the apical surface with 10^5^ PFU IAV Eng195. Inoculum was removed and cells washed three times with PBS. PBS or 0.01 mg/mL ESE treatment occurred at indicated time points. At 24 and 48 hours, apical surfaces were washed with PBS for 30 minutes and virus titre calculated by plaque assay.

### SDS-PAGE analysis of ESE treated IAV

Gradient purified PR8 or concentrations of BSA were mixed with PBS or ESE to a final concentration of 2.5 mg/mL. The samples were further mixed with an equal volume of 2 × SDS sample buffer (0.125 M Tris-HCl [pH 6.8], 4% sodium dodecyl sulphate (SDS), 20% glycerol, 0.004% of bromophenol blue, 10% β-mercaptoethanol) and kept at 95°C for 5 min. Proteins were analysed by 12% SDS-polyacrylamide gel electrophoresis (PAGE) prior to Coomassie R-250 or silver staining (Thermofisher).

### Virus labelling and FACS analysis of virus binding

Gradient purified IAV PR8 was labelled as previously described [26]. Briefly, virus was labelled using a Alexa-Fluor 488 fluorophore labelling kit (Thermofisher, A10235) and unreactive dye was removed by purification through a sucrose cushion. Infectious virus (PR8-488) was determined by plaque assay. For FACS analysis, 2×10^5^ MDCK cells were used per sample. Cells were detached with Trypsin-EDTA and chilled to 4 degrees. PR8-488 was incubated with ESE or infection media for one hour prior to incubation with MDCK cells for one hour on ice. Unbound virus was removed by three washes with PBS and cells fixed with 4% PFA prior to FACS analysis using a BD FACSymphony A1 Flow Cytometer. Single cell populations were gated based on there forward and side scatter using FlowJo software.

### Hemagglutination assays

ESE was serially diluted two-fold in PBS. X31 or PR8 was also serially diluted two-fold and incubated 1:1 for 30 minutes with ESE in 96 well round bottomed plates. Following incubation, 0.5% chicken red blood cells were added to each well and the HA titre calculated after one hour.

### Transmission electron microscopy (TEM)

The negative staining technique was applied. For this approach, purified PR8 was mixed with PBS or 2.5 mg/mL ESE (final concentration) for 30 min at room temperature, then fixed in 2% paraformaldehyde (PFA). A Formvar-coated EM grid was placed, with the Formvar side down, on a drop of virus solution for 1-3 min. The grid was removed, dabbed with filter paper and placed onto a drop of 2.0% phosphotungstic acid (PTA), pH 7.0, for 1 min. Subsequently, the specimen was dried and viewed under a Philips CM10 (FEI, Oregon, USA), operated with a Gatan Orius Sc1000 digital camera (Gatan Microscopical Suite, Digital Micrograph, Version 3.30.2016.0).

### Histological and immunohistological examination

For the histological examination, tissue samples were trimmed after 48 h of formalin fixation and subsequent storing in 70% ethanol until processing, and routinely paraffin wax embedded. Section (2-4 µm) from the tissues in the tolerability study and from left lungs in the infection experiment were prepared and routinely stained with haematoxylin eosin (HE) for histological examination. Consecutive sections from the lungs in the infection experiment were subjected to immunohistological staining for IAV antigen, using the horseradish peroxidase method, a goat anti-IAV (H1N1; virions) antibody (Meridian Life Sciences Inc., Memphis, USA) and a previously published protocol [27].

### Statistical analysis

Data was analysed using Prism package (version 10.1.1). p values were set at 95% confidence interval. Statistical tests used are stated in figure legends. All differences not specifically stated to be significant were not significant (p > 0.05).

## Acknowledgements

This work was funded by the Biotechnology and Biological Sciences Research Council (BBSRC). The authors gratefully acknowledge the Centre for Cell Imaging, University of Liverpool, for their support and assistance in this work.

