## Supplementary files for "Phlorotannin rich *Ascophyllum nodosum* seaweed extract inhibits influenza infection"

**Supplementary material**

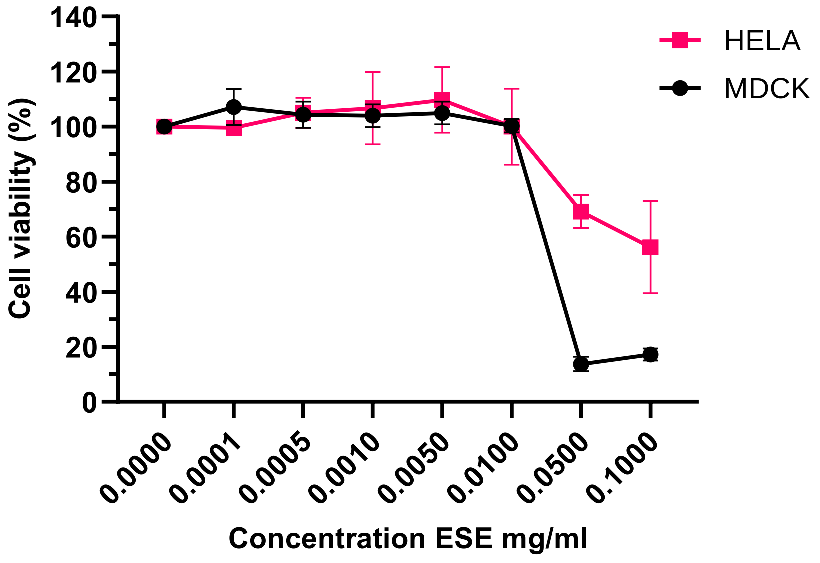

**Supplementary Figure S1.** Cytotoxicity of ESE on MDCK and Hela cells analysed my MTS assay. Cell viability given as a proportion of untreated control. Cell viability below 90% was determined as displaying cytotoxicity. Data represented as mean ± SEM of three independent experiments.

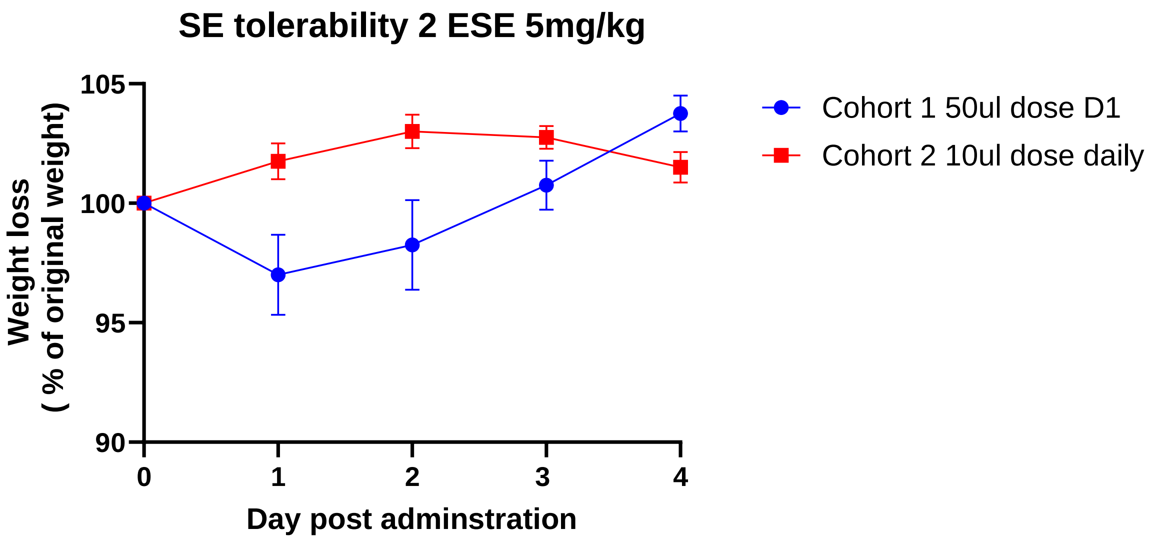

**Supplementary Figure S2**. Tolerability study weight loss to determine the potential pathological effects of ESE in mice. Female C57Bl/6 mice, aged 6-8 weeks, were treated with ESE at a dose of 5 mg/kg either once (in 50 µl PBS) or daily (in 10 µl PBS) for 5 consecutive days. Weight curves. Data represented as mean ± SEM (n=4).

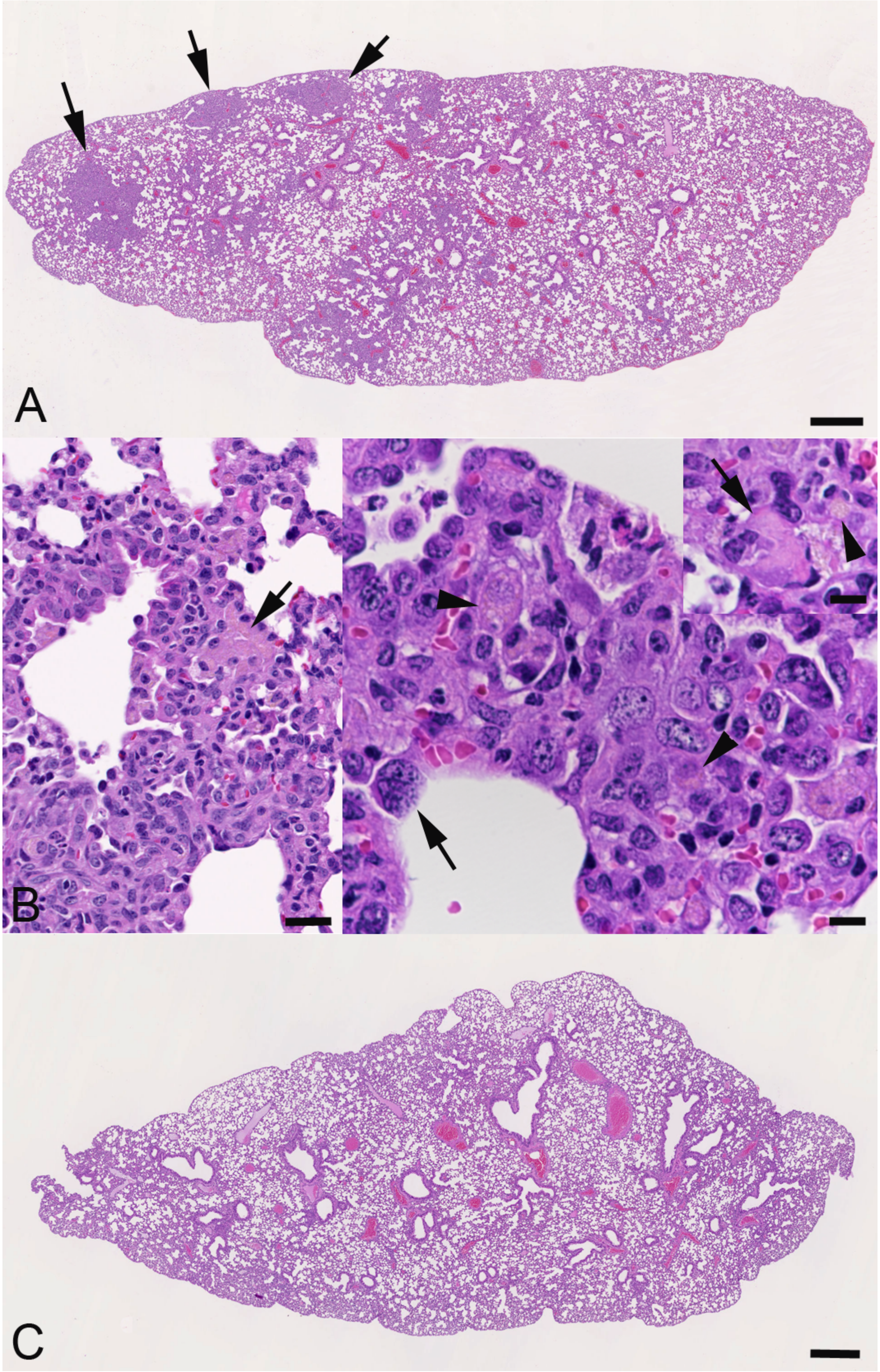

**Supplementary Figure S3**. Tolerability study histological changes to determine the potential pathological effects of ESE in mice. Female C57Bl/6 mice, aged 6-8 weeks, were treated with ESE at a dose of 5 mg/kg either once (in 50 µl PBS) or daily (in 10 µl PBS) for 4 consecutive days. Histological features in the lungs (detailed descriptions are provided in Supplementary Table S1). **A, B**. Animals treated with one dose in 50 µl PBS. **A.** Animal 1.1. There are several random granulomatous infiltrates (arrows). **B.** Closer view of granulomatous infiltrates, with focal aggregate of macrophages containing amorphous eosinophilic and yellowish material (left image: arrow; animal 1.3) and individual vacuolated macrophages that contain granular, slightly yellowish material (right image incl. inset: arrowheads) and that are partly oliogonucleated (arrows) (animal 1.4). **C.** Animal treated with 4 daily doses in 10 µl PBS (animal 2.3). The lung parenchyma is unaltered. HE stain, bars = 500 µm (A, C), 50 µm (B: left image) and 10 µm (B: right image and inset).

**Supplementary table S1:** Weight loss in IAV infected mice treated with ESE, statistical difference. Data was compared using a repeated measures two-way ANOVA (Bonferroni post-test).

| Day 1 | Summary | Adjusted P Value |
|---|---|---|
| Vehicle vs. Prophylactic | **** | <0.0001 |
| Vehicle vs. Time of Infection | ns | 0.4247 |
| Vehicle vs. Therapeutic | * | 0.0164 |
| Day 2 | | |
| Vehicle vs. Prophylactic | ** | 0.0014 |
| Vehicle vs. Time of Infection | ns | 0.0825 |
| Vehicle vs. Therapeutic | * | 0.0173 |
| Day 3 | | |
| Vehicle vs. Prophylactic | ns | 0.1056 |
| Vehicle vs. Time of Infection | ns | >0.9999 |
| Vehicle vs. Therapeutic | ns | 0.9933 |
| Day 4 | | |
| Vehicle vs. Prophylactic | ns | 0.3499 |
| Vehicle vs. Time of Infection | ns | 0.9685 |
| Vehicle vs. Therapeutic | ns | >0.9999 |
| Day 5 | | |
| Vehicle vs. Prophylactic | ns | >0.9999 |
| Vehicle vs. Time of Infection | ns | 0.6966 |
| Vehicle vs. Therapeutic | ns | >0.9999 |

**Supplementary Table S2**. Tolerability study to determine the potential pathological effects of ESE in mice. Female C57Bl/6 mice, aged 6-8 weeks, were treated with ESE at a dose of 5 mg/kg either once (in 50 µl PBS) or daily (in 10 µl PBS) for 5 consecutive days, and culled on day 5. A full histological examination was undertaken on all major tissues/organs.

| **Animal, treatment** | **Histological findings** |
|---|---|
| 1,1 [1 x 50 µl] | **Brain, C1, eyes**: NHAIR |
| | **Heart**: NHAIR |
| | **Respiratory organs:** |

| | |
|---|---|
| | **Trachea**: NHAIR<br>**Lung**: focal areas with a few large alveolar macrophages in lumen and activated type II pneumocytes; focal loose granulomatous infiltrates (macrophages; also in alveolar walls); a few pv leukocyte aggregates (NL, macrophages, LC) |
| | **Alimentary tract:**<br>**Salivary glands, tongue, oesophagus, stomach, SI, LI**: NHAIR<br>**Liver, pancreas**: NHAIR |
| | **Liver, pancreas**: NHAIR |
| | **Urinary tract (kidneys)**: NHAIR |
| | **Endocrine system (pituitary gland, thyroid glands, adrenal glands)**: NHAIR |
| | **Reproductory organs (uterus, ovaries)**: NHAIR |
| | **Haemolymphatic tissues:**<br>**Spleen**: mod sized primary/sec follicles and T cell zones, cell rich red pulp; **MLN, BLN, mand LN**: indistinct follicles and T cell zones, mod cellulatity; **thymus**: NHAIR; **BM**: cell rich, high haematopoietic activity |
| | **Skeletal muscles**: NHAIR |
| 1,2<br>[1 x 50 µl] | **Brain, eyes**: NHAIR |
| | **Heart**: NHAIR |
| | **Respiratory organs:**<br>**Lung**: a few small focal (peribronchial) granulomatous infiltrates; a few pv leukocyte aggregates (NL, macrophages, LC) |
| | **Alimentary tract:**<br>**Salivary glands, tongue, oesophagus, stomach, SI, LI**: NHAIR<br>**Liver, pancreas**: NHAIR |
| | **Urinary tract (kidneys, urinary bladder)**: NHAIR |
| | **Endocrine system (adrenal glands)**: NHAIR |
| | **Reproductory organs (uterus, ovaries)**: NHAIR |
| | **Haemolymphatic tissues:**<br>**Spleen**: mod sized primary/sec follicles and T cell zones, cell rich red pulp; **MLN, BLN**: indistinct follicles and T cell zones, mod cellulatity; **thymus**: NHAIR; **BM**: cell rich, high haematopoietic activity |
| | **Skeletal muscles**: NHAIR |
| | **Skin**: NHAIR |
| 1,3<br>[1 x 50 µl] | **Brain, eyes**: NHAIR |
| | **Heart**: NHAIR |
| | **Respiratory organs:**<br>**Trachea**: NHAIR<br>**Lung**: a few small focal (peribronchial) granulomatous infiltrates; a few pv leukocyte aggregates (NL, macrophages, LC) |
| | **Alimentary tract:**<br>**Salivary glands, tongue, oesophagus, stomach, SI, LI**: NHAIR<br>**Liver, pancreas**: NHAIR |
| | **Liver, pancreas**: NHAIR |
| | **Urinary tract (kidneys, urinary bladder)**: NHAIR |
| | **Endocrine system (pituitary gland, thyroid glands, adrenal glands)**: NHAIR |
| | **Reproductory organs (uterus, ovaries)**: NHAIR |
| | **Haemolymphatic tissues:**<br>**Spleen**: mod sized primary/sec follicles and T cell zones, cell rich red pulp; **BLN, mand LN**: no distinct follicles, mod cellularity; **thymus**: NHAIR; **BM**: cell rich, high haematopoietic activity |
| | **Skeletal muscles**: NHAIR |
| 1,4<br>[1 x 50 µl] | **Brain, C1, eyes**: NHAIR |
| | **Heart**: NHAIR |
| | **Respiratory organs:**<br>**Trachea**: NHAIR<br>**Lung**: multifocal granulomatous infiltrates (macrophages; also in alveolar walls), in larger lesions with embedded NL aggregates; a few pv leukocyte aggregates (NL, macrophages, LC) |
| | **Alimentary tract:** |

| | |
|---|---|
| | **Salivary glands, oesophagus, stomach, SI, LI**: NHAIR<br>**Liver, pancreas**: NHAIR |
| | **Urinary tract (kidneys)**: NHAIR |
| | **Endocrine system (pituitary gland, adrenal glands)**: NHAIR |
| | **Pituitary gland, adrenal glands**: NHAIR |
| | **Reproductory organs (uterus, ovaries)**: NHAIR |
| | **Haemolymphatic tissues:**<br>**Spleen**: mod sized primary/sec follicles and T cell zones, cell rich red pulp; **MLN, BLN**: indistinct follicles and T cell zones, mod cellulatity; **thymus**: NHAIR; **BM**: cell rich, high haematopoietic activity |
| | **Skeletal muscles, femorotibial joint**: NHAIR |
| | **Skin**: NHAIR |
| 2,1<br>[4 x 10 µ] | **Brain, C1, eyes**: NHAIR |
| | **Heart**: NHAIR |
| | **Respiratory organs:**<br>**Trachea**: NHAIR<br>**Lung**: mild focal pv mixed cellular infiltration |
| | **Alimentary tract:**<br>**Salivary glands, oesophagus, stomach, SI, LI**: NHAIR<br>**Liver, pancreas**: NHAIR |
| | **Urinary tract (kidneys, urinary bladder)**: NHAIR |
| | **Endocrine system (pituitary gland, adrenal glands)**: NHAIR |
| | **Reproductory organs (uterus, ovaries)**: NHAIR |
| | **Haemolymphatic tissues:**<br>**Spleen**: mod sized primary/sec follicles and T cell zones, cell rich red pulp; **MLN, mand LN**: indistinct follicles and T cell zones, mod cellularity; **thymus**: NHAIR; **BM**: cell rich, high haematopoietic activity |
| | **Skeletal muscles, femorotibial joint**: NHAIR |
| | **Skin**: NHAIR |
| 2,2<br>[4 x 10 µ] | **Brain, C1, eyes**: NHAIR |
| | **Heart**: NHAIR |
| | **Respiratory organs:**<br>**Trachea**: NHAIR<br>**Lung**: one small artery with mild focal leukocyte rolling and pv accumulation |
| | **Alimentary tract:**<br>**Salivary glands, oesophagus, stomach, SI, LI**: NHAIR<br>**Liver, pancreas**: NHAIR |
| | **Urinary tract (kidneys)**: mild focal mononuclear interstitial infiltration in one kidney |
| | **Endocrine system (pituitary gland, thyroid glands, adrenal glands)**: NHAIR |
| | **Reproductory organs (uterus, ovaries)**: NHAIR |
| | **Haemolymphatic tissues:**<br>**Spleen**: mod sized primary/sec follicles and T cell zones, cell rich red pulp; **MLN**: small section; **thymus**: NHAIR; **BM**: cell rich, high haematopoietic activity |
| | **Skeletal muscles, femorotibial joint**: NHAIR |
| | **Skin**: NHAIR |
| 2,3<br>[4 x 10 µ] | **Brain, C1, eyes**: NHAIR |
| | **Heart**: NHAIR |
| | **Respiratory organs (trachea, lung)**: NHAIR |
| | **Alimentary tract:**<br>**Salivary glands, tongue, oesophagus, stomach, SI, LI**: NHAIR<br>**Liver, pancreas**: NHAIR |
| | **Urinary tract (kidneys, urinary bladder)**: NHAIR |
| | **Endocrine system (adrenal glands)**: NHAIR |
| | **Reproductory organs (uterus, ovaries)**: NHAIR |
| | **Haemolymphatic tissues:** |

| | |
|---|---|
| | **Spleen**: mod sized primary/sec follicles and T cell zones, cell rich red pulp; **MLN, mand LN**: indistinct follicles and T cell zones, mod cellulatity; **thymus**: NHAIR; **BM**: cell rich, high haematopoietic activity |
| | **Skeletal muscles**: NHAIR |
| | **Skin**: NHAIR |
| 2,4 [4 x 10 µ] | **Brain, C1, eyes**: NHAIR |
| | **Heart**: NHAIR |
| | **Respiratory organs (trachea, lung)**: NHAIR |
| | **Alimentary tract:** **Salivary glands, tongue, oesophagus, stomach, SI, LI**: NHAIR **Liver, pancreas**: NHAIR |
| | **Urinary tract (kidneys)**: NHAIR |
| | **Endocrine system (pituitary gland, thyroid glands, adrenal glands)**: NHAIR |
| | **Reproductory organs (uterus, ovaries)**: NHAIR |
| | **Haemolymphatic tissues:** **Spleen**: mod sized primary/sec follicles and T cell zones, cell rich red pulp; **MLN**: indistinct follicles and T cell zones, mod cellulatity; **thymus**: NHAIR; **BM**: cell rich, high haematopoietic activity |
| | **Skeletal muscles, femorotibial joint**: NHAIR |
| | **Skin**: NHAIR |

**Legend**: BLN – bronchial lymph node; C1 – spinal cord at level of C1; LC – lymphocytes; LI – large intestine; mand LN – mandibular lymph node; MLN – mesenteric lymph node; mod – moderate(ly); NHAIR – no histological abnormality is recognised; NL – neutrophilic leukocytes (neutrophils); pv - perivascular; SI – small intestine

**Supplementary Table S3**. Study on the effect of ESE on IAV infection of mice. Female C57Bl/6 mice, aged 6-8 weeks, were infected with a sublethal dose of IAV (cohorts 1-4) and treated with PBS at 3 hpi, then daily (animals 1.1 to 1.6) or with ESE at a dose of 5 mg/kg in 10 µl PBS at 2 h pre-infection, 3 hpi, then daily (animals 2-1 to 2-6), at time of infection, 3 hpi, then daily (animals 3.1 to 3.5) or at 3 hpi, then daily (animals 4.1 to 4.6). Mock-infected mice that received PBS at 3 hpi, then daily (animals 5.1 to 5.3) served as controls. All mice were euthanised at day 5 post infection. A histological examination and immunohistology for IAV antigen was undertaken on the lung.

| Animal No | Histological findings, immunohistology for viral antigen |
|---|---|
| 1.1 | **Lung**: many bronchioles with partly flattened, partly necrotic BEC, partly with individual necrotic bronchial/-iolar BEC, with LC dominated bronchial and pb infiltration, abundant degenerate cells, debris and NL in lumen; adjacent focal parenchymal areas with desquamation of AM/type II pn, sometimes necrotic cells, activated type II pn, some NL and LC; vasculitis and pv LC-dominated mononuclear infiltration **vAg**: numerous bronchioles with mod number of pos BEC (intact and degen, individual or patches; in lumen) and AEC (and macrophages) in parenchymal infiltrates adjacent to affected bronchioles |
| 1.2 | **Lung**: bronchus with focal area of BEC necrosis and LC dominated bronchial and pb infiltration, degenerate cells, debris and NL in lumen; two small focal |

| | |
|---|---|
| | parenchymal areas with desquamation of AM/type II pn, activated type II pn, some NL and LC<br>**vAg**: bronchus with large patch of pos BEC (intact and degen, individual or patches; in lumen), one bronchiole with a few individual pos BEC; AEC (and macrophages) in parenchymal infiltrates |
| 1.3 | **Lung**: bronchus with focal BEC necrosis, loss and debris in lumen, with LC dominated bronchial and pb infiltration; focal parenchymal area with desquamation of AM/type II pn, some necrotic cells, activated type II pn, some NL and LC; mild vasculitis and pv LC-dominated mononuclear infiltration<br>**vAg**: bronchus with almost diffuse pos BEC; several bronchioles with large patches of pos BEC; AEC (and macrophages) mainly in parenchymal infiltrate |
| 1.4 | **Lung**: one large parenchymal area with embedded bronchiole exhibiting BEC necrosis, loss and debris in lumen, with LC dominated bronchial and pb infiltration; parenchyma with desquamation of AM/type II pn, some necrotic cells, activated type II pn, some NL and LC; vasculitis and pv LC-dominated mononuclear infiltration<br>**vAg**: in one large area bronchioles with variable mount of pos BEC (intact and degen); pos AEC (and macrophages) mainly in extensive parenchymal infiltrate |
| 1.5 | **Lung**: one large parenchymal area with embedded bronchiole exhibiting complete BEC necrosis, loss and debris in lumen, with LC dominated bronchial and pb infiltration; parenchyma with desquamation of AM/type II pn, some necrotic cells, activated type II pn, some NL and LC; vasculitis and pv LC-dominated mononuclear infiltration<br>**vAg**: in one large area bronchioles with variable amount of pos BEC (intact and degen); pos AEC (and macrophages) mainly in extensive parenchymal infiltrate |
| 1.6 | **Lung**: bronchus and several bronchioles with (partly) necrotic BEC, with LC dominated bronchial and pb infiltration, abundant degen cells, debris and NL in lumen; adjacent focal parenchymal areas with desquamation of AM/type II pn, some necrotic cells, activated type II pn, some NL and LC; mild vasculitis and pv LC-dominated mononuclear infiltration<br>**vAg**: several bronchioles with mod number of pos BEC (intact and degen, individual or patches; in lumen); AEC (and macrophages) in parenchymal infiltrates adjacent to affected bronchioles |
| 2.1 | **Lung**: a few bronchioles with complete necrosis and loss of BEC, filled with abundant degen cells, debris and proteinaceous material; with LC dominated bronchial and pb infiltration; one to several adjacent focal granulomatous infiltrates with embedded NL (similar also in alveoli); also small parenchymal areas with desquamation of AM/type II pn, some necrotic cells, activated type II pn, some NL and LC; mild vasculitis and pv LC-dominated mononuclear infiltration<br>**vAg**: in BEC and cell free in material in lumen of bronchi |
| 2.2 | **Lung**: two larger and a few small nodular focal pb granulomatous infiltrates with embedded individual NL and aggregates of LC; large bronchiole with activated, partly hyperplastic BEC and attached clumped proteinaceous material<br>**vAg**: individual pos BEC in several bronchioles, also cell free in material in lumen of bronchi |
| 2.3 | **Lung**: NHAIR<br>**vAg**: one bronchiole with one pos BEC; one small area with a few pos AEC |
| 2.4 | **Lung**: mod focal pv LC-dominated mixed cellular infiltration and adjacent parenchymal area with proteinaceous material and degen cells in alveolar lumina and mixed inflammatory infiltrate; small focal granulomatous parenchymal infiltrate adjacent to small bronchiole<br>**vAg**: neg |
| 2.5 | **Lung**: one large bronchiole with focal pb LC dominated infiltration and adjacent focal consolidated area with desquamation of AM/type II pn, some necrotic cells, activated type II pn, some NL and LC<br>**vAg**: one bronchiole with one pos BEC; focal lesion with numerous pos AEC (and macrophages) within and immediately adjacent to looser consolidated areas |

| | |
|---|---|
| 2.6 | **Lung**: mild multifocal pv LC-dominated infiltration, one pv parenchymal area with degen cells in alveolar lumina and mixed inflammatory infiltrate; one bronchiole with proteinaceous material and degen cells in lumen<br>**vAg**: one bronchiole with a patch of pos BEC, also cell free in material in lumen of bronchi |
| 3.1 | **Lung**: NHAIR<br>**vAg**: neg |
| 3.3 | **Lung**: mod multifocal pv LC-dominated, partly mixed cellular infiltration; one vessel with marked focal NL infiltration and adjacent parenchymal area with marked NL infiltration; several nodular focal granulomatous parenchymal infiltrates adjacent to bronchioles, partly NL dominated; a few bronchioles with mild to mod BEC hyperplasia and some debris in lumen<br>**vAg**: neg |
| 3.4 | **Lung**: NHAIR<br>**vAg**: neg |
| 3.5 | **Lung**: multifocal (nodular) granulomatous parenchymal infiltrates adjacent to bronchioles, with variable proportions of NL; a few bronchioles with mild BEC hyperplasia and some debris in lumen<br>**vAg**: neg |
| 3.6 | **Lung**: NHAIR, apart from rare very small focal granulamatous infltrates<br>**vAg**: neg |
| 4.1 | **Lung**: focal area with mod pb and pv LC infiltration; focal consolidated area with desquamation of AM/type II pn, some necrotic cells, activated type II pn, some NL and LC<br>**vAg**: one large area with several bronchioles with mod number of pos BEC (intact and degen, individual or patches; in lumen, with some clumped cell free antigen) and AEC (and macrophages) in parenchymal infiltrates adjacent to affected bronchioles |
| 4.2 | **Lung**: NHAIR, apart from very mild focal LC aggregates<br>**vAg**: one bronchiole with one pos BEC |
| 4.3 | **Lung**: focal area with mod pb and pv LC infiltration; adjacent focal area with desquamation of AM/type II pn, some necrotic cells, activated type II pn, some NL and LC, alveolar oedema<br>**vAg**: two bronchioles with large patch of pos BEC; pos AEC and macrophages in alveoli in the periphery of focal parenchymal area with inflammatory infiltration |
| 4.4 | **Lung**: NHAIR<br>**vAg**: neg |
| 4.5 | **Lung**: bronchus with mod to marked pb LC-dominated (and focally NL dominated) infiltration, some degen sloughed off BEC and abundant NL in lumen<br>**vAg**: bronchus with several individual pos BEC and some degen pos BEC in lumen (with debris) |
| 4.6 | **Lung**: focal area with mod pb and pv LC infiltration (with some NL); focal area with desquamation of AM/type II pn, some necrotic cells, activated type II pn, some NL and LC, and leukocyte recruitment<br>**vAg**: focal area with a few pos BEC in some bronchioles and a few adjacent pos AEC (and macrophages), several pos AEC and macrophages in focal parenchymal area with inflammatory infiltration |
| 5.1 | **Lung**: NHAIR; **vAg**: neg |
| 5.2 | **Lung**: NHAIR; **vAg**: neg |
| 5.3 | **Lung**: NHAIR; **vAg**: neg |

**Legend**: AEC – alveolar epithelial cells; AM – alveolar macrophages; BEC – bronchiolar epithelial cells; degen – degenerate; LC – lymphocytes; neg – negative; NHAIR – no histological abnormality is recognised; NL – neutrophilic leukocytes (neutrophils); pn – pneumocytes; pos – positive; pv – perivascular; vAg – viral antigen

Cohort 1: PBS 3 hpi, then daily
Cohort 2: ESE 5mg/kg, 2 h pre-inf, 3 hpi, then daily
Cohort 3: 5mg/kg, time of inf, 3 hpi, then daily
Cohort 4: 5 mg/kg, 3 hpi, then daily
Cohort 5: Uninfected; PBS, 3 hpi, then daily
